# A phospho-switch at Acinus-Serine^437^ controls autophagic responses to Cadmium exposure and neurodegenerative stress

**DOI:** 10.1101/2021.07.27.453988

**Authors:** Nilay Nandi, Zuhair Zaidi, Charles Tracy, Helmut Krämer

## Abstract

Neuronal health depends on quality control functions of autophagy, but mechanisms regulating neuronal autophagy are poorly understood. Previously, we showed that in Drosophila starvation-independent quality control autophagy is regulated by Acinus and the Cdk5-dependent phosphorylation of its serine^437^ (Nandi et al., 2017). Here, we identify the phosphatase that counterbalances this activity and provides for the dynamic nature of Acinus-S437 phosphorylation. A genetic screen identified six phosphatases that genetically interacted with an Acinus gain-of-function model. Among these, loss of function of only one, the PPM-type phosphatase Nil (CG6036), enhanced pS437-Acinus levels. Cdk5-dependent phosphorylation of Acinus serine^437^ in *nil*^1^ animals elevates neuronal autophagy and reduces the accumulation of polyQ proteins in a Drosophila Huntington’s disease model. Consistent with previous findings that Cd^2+^ inhibits PPM-type phosphatases, Cd^2+^-exposure elevated Acinus-serine^437^ phosphorylation which was necessary for increased neuronal autophagy and protection against Cd^2+^-induced cytotoxicity. Together, our data establish the Acinus- S437 phospho-switch as critical integrator of multiple stress signals regulating neuronal autophagy.

## Introduction

A key process for maintaining cellular fitness is autophagy, here short for macroautophagy (Fleming and Rubinsztein, 2020; Menzies et al., 2015). Starvation induces non-selective autophagy which contributes to reclaiming molecular building blocks (Levine and Kroemer, 2019). In neurons and other long-lived cells, quality control of proteins and organelles is an additional critical function of autophagy (Dong et al., 2021; Evans and Holzbaur, 2020; Kroemer et al., 2010). The importance of the quality control function of starvation-independent basal autophagy was demonstrated by mutations in core autophagy components in mice and flies: cell-type specific loss of Atg5 or Atg7 triggers rapid neurodegeneration (Hara et al., 2006; Juhasz et al., 2007; Komatsu et al., 2006) or cardiac hypertrophy (Nakai et al., 2007). Moreover, elevated basal autophagy can successfully reduce the polyQ load in models of Huntington’s disease or spinocerebellar ataxia type 3 (SCA3) and reduce neurodegeneration (Bilen and Bonini, 2007; Jaiswal et al., 2012; Nandi et al., 2014, 2017; Ravikumar et al., 2004). Both modes of autophagy use core autophagy proteins to initiate the generation of isolation membranes (also known as phagophores), promote their growth to autophagosomes, and finally promote their fusion with lysosomes or late endosomes to initiate degradation of captured content (Mizushima, 2017). Although the rapid induction of autophagy in response to nutrient deprivation is well described (Galluzzi et al., 2017), much less is known about the modulation of basal levels of autophagy in response to cellular stress.

We previously identified Acinus (Acn) as a regulator of starvation-independent quality control autophagy in Drosophila (Haberman et al., 2010; Nandi et al., 2014, 2017). Acn is a conserved protein enriched in the nucleus and, together with Sap18 and RNPS1, forms the ASAP complex (Murachelli et al., 2012; Schwerk et al., 2003). The ASAP complex can regulate alternative splicing by interacting with the exon junction complex, spliceosomes and messenger ribonucleoprotein particles (Hayashi et al., 2014; Malone et al., 2014; Tang et al., 1995). In mammals and Drosophila, Acn levels are regulated by its Akt1-dependent phosphorylation which inhibits caspase-mediated cleavage (Hu et al., 2005; Nandi et al., 2014). Furthermore, Acn stability is enhanced by Cdk5-mediated phosphorylation of the conserved Serine-437. Acn levels are elevated by the phospho-mimetic Acn^S437D^ mutation and reduced by the phospho-inert Acn^S437A^ (Nandi et al., 2017). The stress-responsive Cdk5/p35 kinase complex (Su and Tsai, 2011) regulates multiple neuronal functions including synapse homeostasis and axonal transport (Klinman and Holzbaur, 2015; Lai and Ip, 2015; McLinden et al., 2012), in addition to its role in autophagy (Nandi and Krämer, 2018; Shukla and Giniger, 2019). Phosphorylation-induced stabilization of Acn increases basal, starvation-independent autophagy with beneficial consequences including reduced polyQ load in a Drosophila Huntington’s disease model and prolonged life span (Nandi et al., 2014, 2017). The detailed mechanism by which Acn regulates autophagy is not well understood, but is likely to involve the activation of Atg1 kinase activity as autophagy-related and unrelated functions of Atg1 are enhanced by elevated Acn levels (Nandi et al., 2014, 2017; Tyra et al., 2020). Identification of Acn in a high-content RNAi screen for genes promoting viral autophagy in mammals (Orvedahl et al., 2011) suggests a conserved role in regulating starvation-independent autophagy.

Detailed examination of the cell type-specific changes in levels and phosphorylation of Acn in photoreceptor neurons of developing larval eye discs revealed a highly dynamic pattern (Nandi et al., 2014, 2017). This motivated us to investigate the role of serine-threonine phosphatases in counteracting Cdk5/p35 kinase-mediated Acn phosphorylation. In a targeted screen, we identified CG6036, a member of the PPM family of protein phosphatases as critical for controling the phospho-switch on Acn-S437. PPM-type phosphatases are dependent on Mg^2+^ or Mn^2+^ as co-factor for their activity (Kamada et al., 2020). They are not inhibited by the broad-spectrum phosphatase inhibitor okadaic acid, in contrast to PPP-type phosphatases and do not require the regulatory subunits characteristic for PPP-type phosphatases. Instead, the PPM family contains additional domains and conserved motifs, which can determine its substrate specificity (Andreeva and Kutuzov, 2001; Shi, 2009; Tong et al., 1998). Intriguingly, PPM family members play a pivotal role in different physiological or pathological processes that are responsive to cellular stress signaling, including regulation of AMPK (Davies et al., 1995), Tak1 (Hanada et al., 2001), or other MAP kinases (Hanada et al., 1998; Maeda et al., 1994; Shiozaki et al., 1994; Takekawa et al., 1998).

Our findings indicate that the CG6036 phosphatase, through its effect on Acn phosphorylation, regulates neuronal responses to proteostasis or toxicological stress. We renamed this phosphatase “Nilkantha” (Nil) and from here on will refer to the CG6036 phosphatase as Nil.

## Results

### Nil phosphatase regulates phosphorylation of the conserved serine 437 of Acn

To identify phosphatases responsible for modulating Acn function, we performed a targeted RNAi screen of the 37 non CTD-type serine-threonine phosphatases encoded in the Drosophila genome (Supplemental Table 1). To test the effect of these phosphatases on Acn function, we used an eye-specific sensitized genetic system. GMR-Gal4-driven expression of UAS-Acn^WT^ at 28°C yields a rough-eye phenotype (Figure 1A), that is modified by genetic enhancers or suppressors (Nandi et al., 2014, 2017). We reasoned that knocking down a phosphatase responsible for dephosphorylating Acn would elevate the levels of phosphorylated Acn and hence stabilize the Acn protein, resulting in an enhancement of the eye roughness induced by UAS-Acn^WT^. Among the serine-threonine phosphatases encoded by the Drosophila genome, RNAi lines targeting *CG6036*, *CG15035*, *PpD6*, *Pp1-13C*, *flapwing (flw)* and *microtubule star (mts)* exhibited enhancement of Acn-induced eye roughness yielding a severely rough and reduced eye (Figure 1A-G, Supplemental Table 1, Figure 1-figure supplement 1). By contrast, expression of these RNAi transgenes by themselves did not result in visible eye phenotypes (Figure 1H-N, Supplemental Table 1).

**Figure 1.**
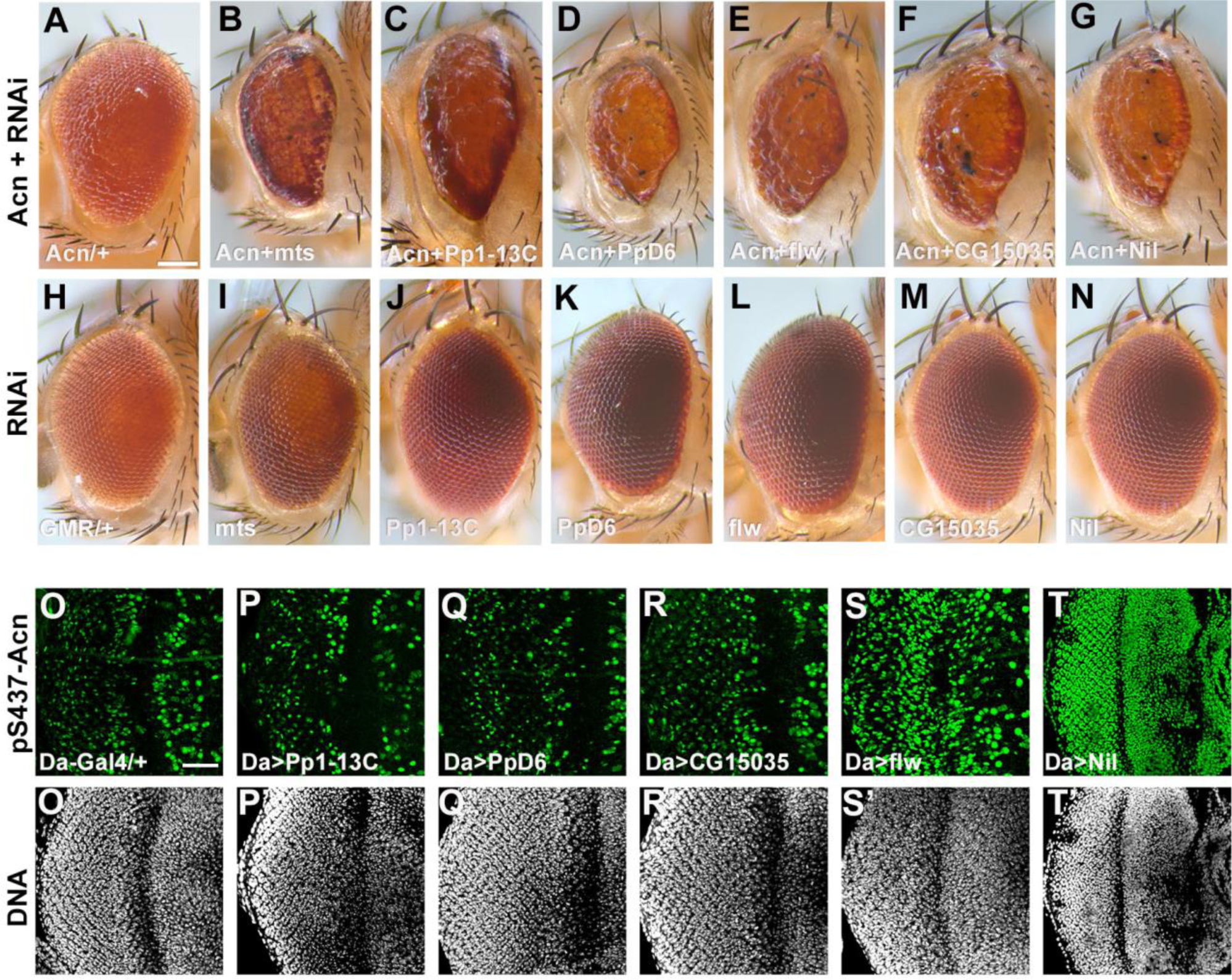
A genetic screen identifies Nil as the Acinus-S437 phosphatase. (A–N) Micrographs of eyes in which GMR-Gal4 drives expression of Acn^WT^ (A), Acn^WT^ + mts–RNAi (B), Acn^WT^ + Pp1-13C RNAi (C), Acn^WT^ + PpD6 RNAi (D), Acn^WT^ + flw RNAi (E), Acn^WT^ + CG15035 RNAi (F), Acn^WT^ + nil RNAi (G), mts–RNAi (I), Pp1-13C–RNAi (J), PpD6–RNAi (K), flw–RNAi (L), CG15035–RNAi (M), nil–RNAi (N) and H represents GMR-Gal4 control. (O–T) Projection of confocal micrographs of larval eye discs stained for pS437-Acn (green) and DNA from Da-Gal4 (O, O’), Da-Gal4, Pp1-13C RNAi (P, P’), Da-Gal4, PpD6 RNAi (Q, Q’), Da-Gal4, flw RNAi (R, R’), Da-Gal4, CG15035 RNAi (S, S’), Da-Gal4, nil RNAi (T, T’). Scale bar in A is 100 µm for A-N, scale bar in O is 40 µm for O-T. Genotypes are listed in Supplemental Table 3.

To test whether these genetic interactions reflect direct effects on the phosphorylation status of Acn, we used a phospho-specific antibody raised against pS437-Acn (Nandi et al., 2017) to stain eye discs in which these phosphatases had been knocked down using the ubiquitously expressed Da-Gal4 driver. No change in Acn phosphorylation resulted from knockdown of the phosphatases Pp1-13C, PpD6, or CG15035 (Figure 1O-R). By contrast, eye discs with *flw* knockdown displayed an altered pattern of pS437-Acn positive cells (Figure 1S). Moreover, knocking down *mts* with Da-Gal4 resulted in larval lethality.Interestingly, the PP2A phosphatase Mts, a member of the STRIPAK complex, and the PP1 phosphatase Flw regulate upstream components of Hippo/Yorkie signaling (Gil-Ranedo et al., 2019; Neal et al., 2020; Ribeiro et al., 2010; Yang et al., 2012). Furthermore, Yorkie’s growth promoting activity is regulated by Acn activity (Tyra et al., 2020). Taken together, this suggests that the strong genetic interactions of Acn with the Mts and Flw phosphatases might reflect additive effects on Yorkie activity rather than direct effects on Acn phosphorylation.

By contrast, knock down of Nil (CG6036) yielded a dramatic enhancement of Acn phosphorylation at serine^437^ compared to wild-type controls (Figure 1O,T). Given this robust increase of Acn phosphorylation, we further explored the role of Nil in regulating Acn function.

Nil is a member of the PPM family of phosphatases characterized by multiple conserved acidic residues (Figure 2A) that contribute to a binuclear metal center critical for phosphatase activity (Das et al., 1996; Pan et al., 2013). To further test the role of the Nil phosphatase in regulating Acn-S437 phosphorylation, we used CRISPR-Cas9 to generate the *nil*^1^ deletion allele that eliminates the majority of the conserved phosphatase domain (Figure 2A). Antennal discs and larval fat bodies from *nil*^1^ wandering larvae displayed a dramatic increase in Acn-S437 phosphorylation compared to wild-type controls (Figure 2B- E). A similar robust enhancement of pS437-Acn staining was seen in *nil*^1^ mutant eye discs compared to the controls (Figure 2F,G). Overexpressing wild-type Nil or the human PPM1B homolog of Nil restored phosphatase activity in *nil*^1^ mutant eye discs (Figure 2H,I). Multiple sequence alignment pointed to aspartate-231 of Nil as an acidic residue critical for metal binding and phosphatase activity (Kamada et al., 2020). Mutation of this aspartate residue to asparagine generated the Nil^D231N^ point mutant; its expression in *nil*^1^ mutant eye discs failed to restore phosphatase activity (Figure 2J). Moreover, the rough-eye phenotype induced by Acn overexpression using the GMR-Gal4 driver was suppressed by co-expression of wild-type Nil, but not Nil^D231N^ (Figure 2-figure supplement 1A-F). Additionally, overexpression of wild-type Nil, but not the inactive Nil^D231N^ mutant, in larval eye discs reduced pS437-Acn levels compared to GMR-Gal4 only controls (Figure 2-figure supplement 1G-I).

**Figure 2.**
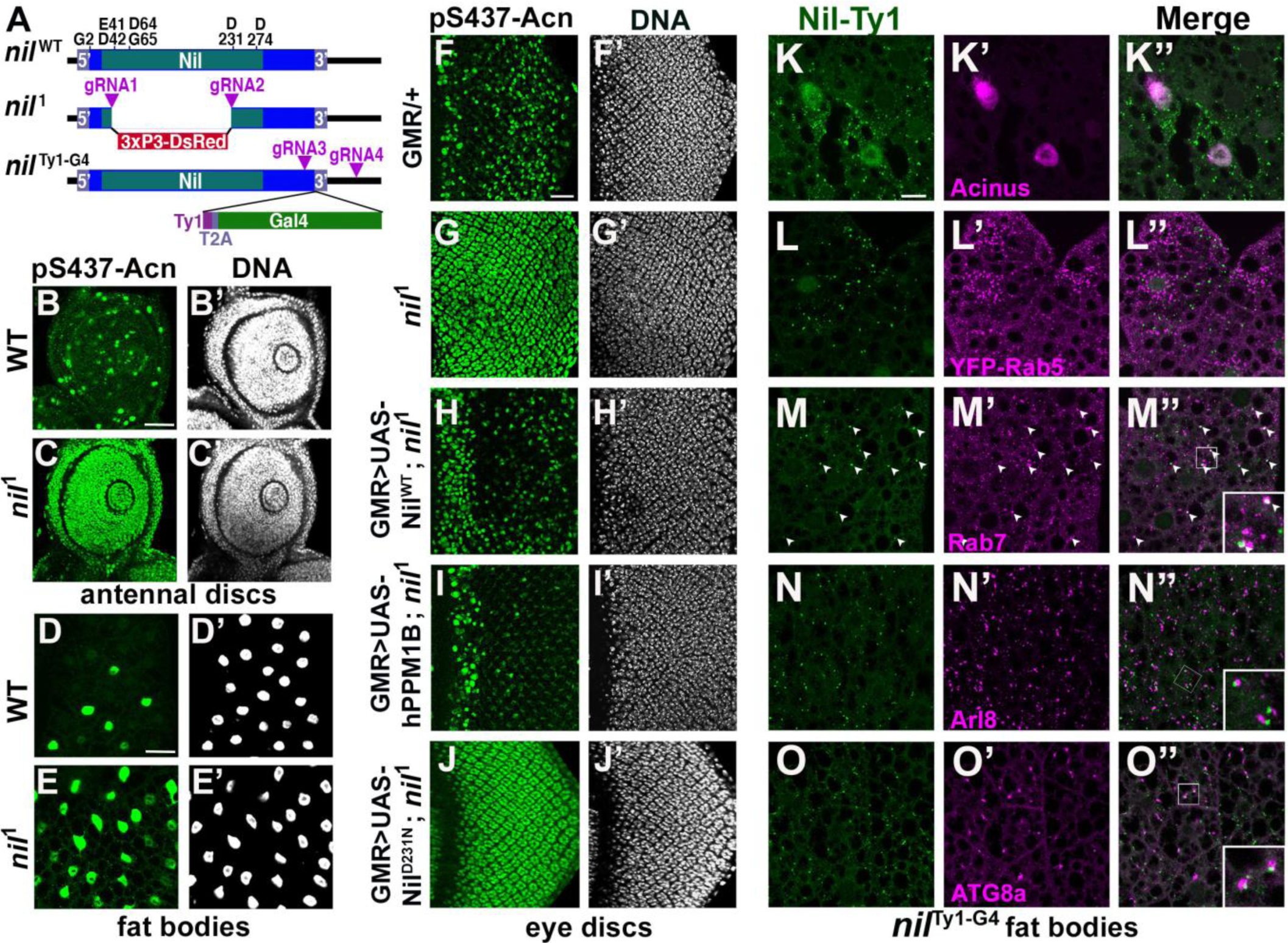
Nil loss and gain-of-function regulates Acinus phosphorylation. (A) Diagram depicting the Nil^WT^ protein with amino acids highly conserved in the PPM family of phosphatases, the *nil*^1^ allele with its 3xP3-DsRed insertion and the multicistronic *nil*^Ty1-G4^ allele with Ty1 tag and T2A-co-expressed Gal4. (B–E) Projection of confocal micrographs of larval antennal discs (B, C) and fat bodies (D, E) stained for pS437-Acn (green) and DNA from *w*^1118^ and *nil*^1^. (F–J) Projection of confocal micrographs of larval eye discs stained for pS437-Acn (green) and DNA from GMR-Gal4/+ (F, F’), *nil*^1^ (G, G’), GMR-Gal4, UAS-Nil^WT^; *nil*^1^ (H, H’), GMR-Gal4, UAS- hPPM1B; *nil*^1^ (I, I’), GMR-Gal4, UAS-Nil^D231N^; *nil*^1^ (J, J’). (K–O) Projection of confocal micrographs of larval fat bodies from *nil*^Ty1-G4^ all stained for Ty1(green) and Acn (magenta) (K- K’’), YFP-Rab5 (magenta) (L-L’’), Rab7 (magenta) (M-M’’), Arl8 (magenta) (N- N’’), or Atg8a (magenta) (O-O’’). Arrowheads in M-M’’ indicate colocalization of Nil-Ty1 with Rab7 in cytosolic punctae. Notice that projections in L to O represent apical regions largely excluding nuclei. Scale bar in B is 40 µm for B-E and in F is 20 µm for F-O. Genotypes are listed in Supplemental Table 3.

### Nil phosphatase localizes to the nucleus and to endo-lysosomal compartments

To gain insights in how this phosphatase can regulate Acn phosphorylation and function we examined its subcellular localization. We generated the *nil*^Ty1-G4^ allele expressing Ty1- tagged Nil phosphatase and Gal4 under control of endogenous nil promoter (Figure 2A, Figure 2-figure supplement 2F). To examine localization of the Nil phosphatase we stained *nil*^Ty1-G4^ animals using antibodies against the Ty1 tag. Consistent with the previously reported low expression levels (Brown et al., 2014), we could barely detect Ty1-tagged Nil phosphatase in eye discs of wandering third instar larvae (Figure 2-figure supplement 2A,B). However, Ty1-tagged Nil phosphatase was abundant in nuclei of third instar larval fat bodies (Figure 2-figure supplement 2C-E), consistent with its role in regulating phosphorylation of the primarily nuclear Acn protein (Haberman et al., 2010; Nandi et al., 2014, 2017). Co-staining of third instar larval fat bodies of *nil*^Ty1-G4^ larvae indicated expression of endogenous Nil phosphatase in Acn-positive nuclei (Figure 2K). Moreover, cytosolic punctae positive for the Nil phosphatase prompted us to examine its possible localization to endo-lysosomal compartments. We compared Nil localization to that of the early endosomal marker YFP- Rab5 (Dunst et al., 2015), but we found no co-localization of Ty1-positive Nil phosphatase punctae with YFP-Rab5 (Figure 2L). We further co-stained *nil*^Ty1-G4^ larval fat bodies with antibodies against Ty1 and Rab7, Arl8 or ATG8a. Rab7 marks late endosomes (Numrich and Ungermann, 2014), Arl8 lysosomes (Rosa-Ferreira et al., 2018) and ATG8a autophagosomes and early autolysosomes (Klionsky et al., 2016). We observed many of the prominent Ty1-stained Nil phosphatase punctae to colocalize with Rab7 (arrowheads in Figure 2M) or to be adjacent to Arl8-marked lysosomes and Atg8a-marked autophagosomes/autolysosomes (Figure 2N,O). This localization suggested a possible involvement of the Nil phosphatase in regulating components of endo-lysosomal or autophagic trafficking (2K-O, Figure 2-figure supplement 2) consistent with our previously described role of Acn in this pathway (Nandi and Krämer, 2018).

### Phosphorylation of Acn Serine 437 in *nil*^1^ animal elevates basal autophagy

Increased Acn-S437 phosphorylation elevates the level of basal, starvation-independent autophagy (Nandi et al., 2017). We therefore tested whether *nil*^1^ null animals exhibited increased levels of autophagy. Consistent with increased autophagy, *nil*^1^ displayed increased ratio of lipidated Atg8a-II to Atg8a-I (Figure 3A,B). Furthermore, we examined endogenous Atg8a in eye and antennal discs of fed wandering third instar larvae. We observed higher numbers of Atg8a-positive punctae in *nil*^1^ imaginal discs compared to wild type, possibly indicating elevated levels of autophagy (Figure 3C-H). Larval fat bodies are a well-established model in Drosophila for investigating autophagy (Rusten et al., 2004; Scott et al., 2004). In fat bodies of fed 96 hr larvae, we observed few Atg8a-positive punctae but their number increased upon a 4-hr amino acid starvation (Figure 3I,M,Q,R). However, numerous ATG8a-positive structures were found in fed *nil*^1^ larval fat bodies (Figure 3J,Q) and their abundance further increased on starvation (Figure 3N,R).

**Figure 3.**
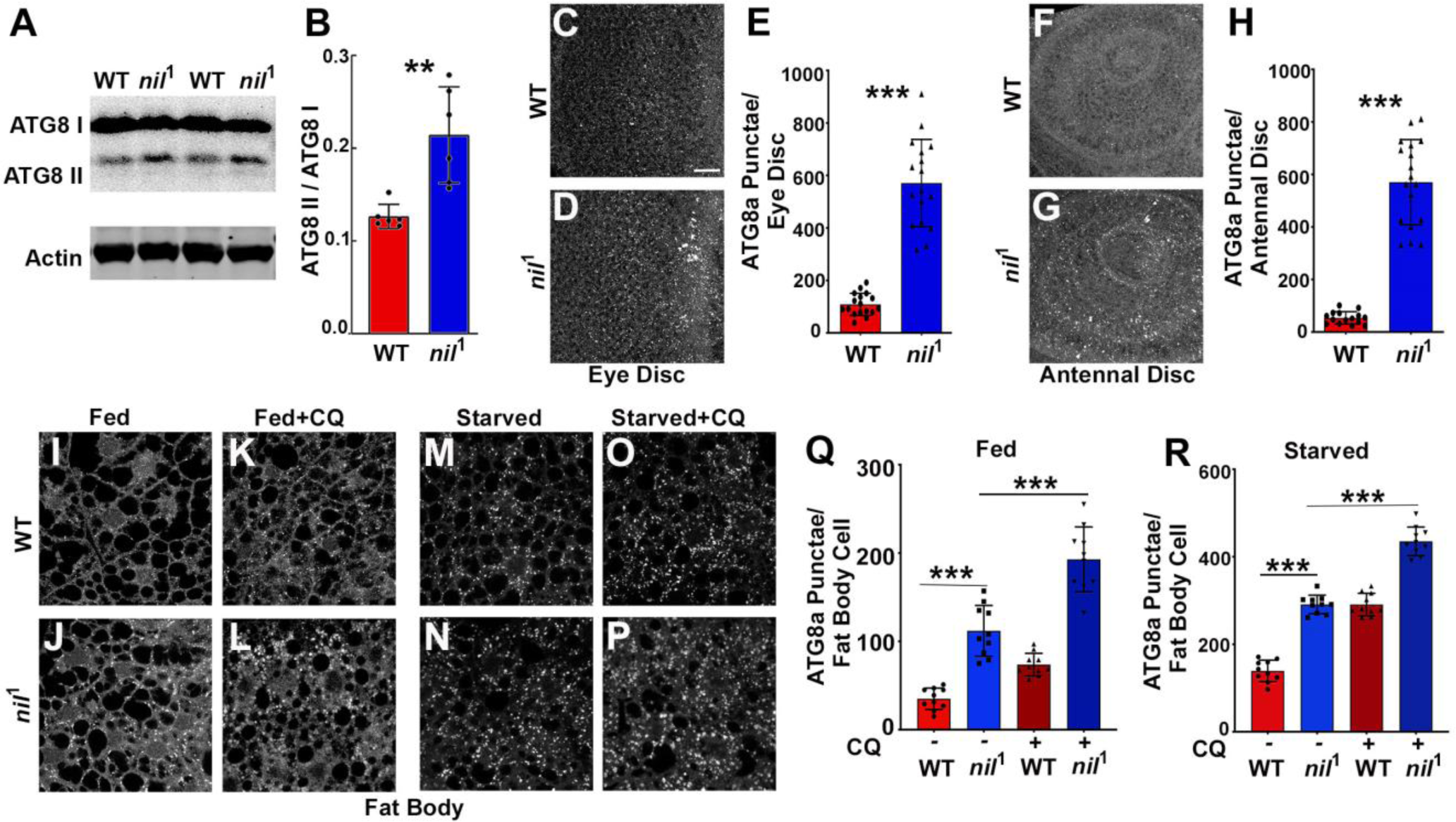
Loss of Nil enhances autophagic flux. (A-B) Western blot of lysates from adult heads of *w*^1118^ and *nil*^1^ probed for ATG8a (A). Quantification of ATG8a-II to ATG8a-I ratio from Western blots as in A. Data are from 6 different lysates from three experimental repeats. Bar graphs show mean ±SD. **p<0.01 (B). (C-E) Projection of confocal micrographs of fed *w*^1118^ and *nil*^1^ larval eye discs (C, D) stained for Atg8a. (E) Quantification of Atg8a punctae in eye discs (C, D) of fed *w*^1118^ and *nil*^1^. Data are from 15 larvae taken from three experimental repeats. Bar graphs show mean ±SD. ***p<0.001. (F-H) Projection of confocal micrographs of fed *w*^1118^ and *nil*^1^ antennal discs (F, G) stained for Atg8a. (H) Quantification of Atg8a punctae in antennal discs (F, G) of fed *w*^1118^ and *nil*^1^. Data are from 15 larvae from three experimental repeats. Bar graphs show mean ±SD. ***p<0.001. (I-P) Projection of confocal micrographs encompassing 6 to 8 cells of *w*^1118^ and *nil*^1^ larval fat bodies aged 96 h after egg laying, either fed (I-L) or amino-acid starved for 4h in 20% sucrose solution (M-P) stained for Atg8a. Larvae were matched for size. To assess autophagic flux, for panels K, L and O, P lysosomal degradation was inhibited with chloroquine (CQ). (Q-R) Quantification of Atg8a punctae in fed (Q) or starved (R) *w*^1118^ and *nil*^1^ larval fat bodies averaged from six to eight cells each. Data are from 10 larvae from three experimental repeats. Bar graphs show mean ±SD. ***p<0.001. Scale bar in C is 20 µm for C-D, F-G, I-P. Genotypes are listed in Supplemental Table 3.

The increased number of ATG8a punctae in *nil*^1^ animals may either represent a failure of autophagosomes to fuse with lysosomes, or an enhanced autophagy induction and flux. To distinguish between these possibilities, we inhibited lysosomal acidification and degradation with chloroquine (Mauvezin et al., 2014). For starved wild-type and *nil*^1^ larval fat bodies, chloroquine treatment further elevated ATG8a staining after starvation, consistent with enhanced autophagy flux in these starved tissues (Figure 3O,P,R). Most importantly, treating fed 96-hr larval fat bodies of *nil*^1^ animals with chloroquine significantly enhanced the number of ATG8a positive punctae demonstrating an elevated autophagic flux (Figure 3K,L,Q). Enhanced autophagic flux in *nil*^1^ animals in which phosphorylation of Acn S437 is robustly elevated is consistent with our previous work, that showed by several methods that phosphorylation of this residue elevates autophagic flux (Nandi et al., 2017).

### Cdk5-p35 kinase complex triggers Acn S437 phosphorylation in *nil*^1^ animal

PPM1-type serine-threonine phosphatases can negatively regulate stress-responsive MAPK kinase cascades by directly dephosphorylating and thereby deactivating MAP kinases or MAPK activating kinases (Hanada et al., 1998; Takekawa et al., 1998). Therefore, we examined a possible role of the Nil phosphatase in regulating the activity of the MAPK cascades and their possible involvement in phosphorylating Acn-S437 in *nil*^1^ mutants. We examined phosphorylation of three MAP Kinase family members: ERK, SAPK/JNK and p38b MAPK in larval eye discs of *nil*^1^ animals. We did not observe changes in phosphorylation of ERK or SAPK/JNK in *nil*^1^ eye-antennal discs compared to wild type (Figure 4A-D). p38bMAPK exhibited elevated phosphorylation in some undifferentiated cells anterior to the morphogenetic furrow, but not in the developing photoreceptor neurons of *nil*^1^ eye disc compared to wild type (Figure 4E-F). Furthermore, we have demonstrated earlier that phosphorylation of Acn-S437 remains unchanged in *p38b* mutant eye discs compared to wild type (Nandi et al., 2017), arguing against an involvement of p38b MAPK in regulating Acn phosphorylation at serine 437 in *nil*^1^ animals.

**Figure 4.**
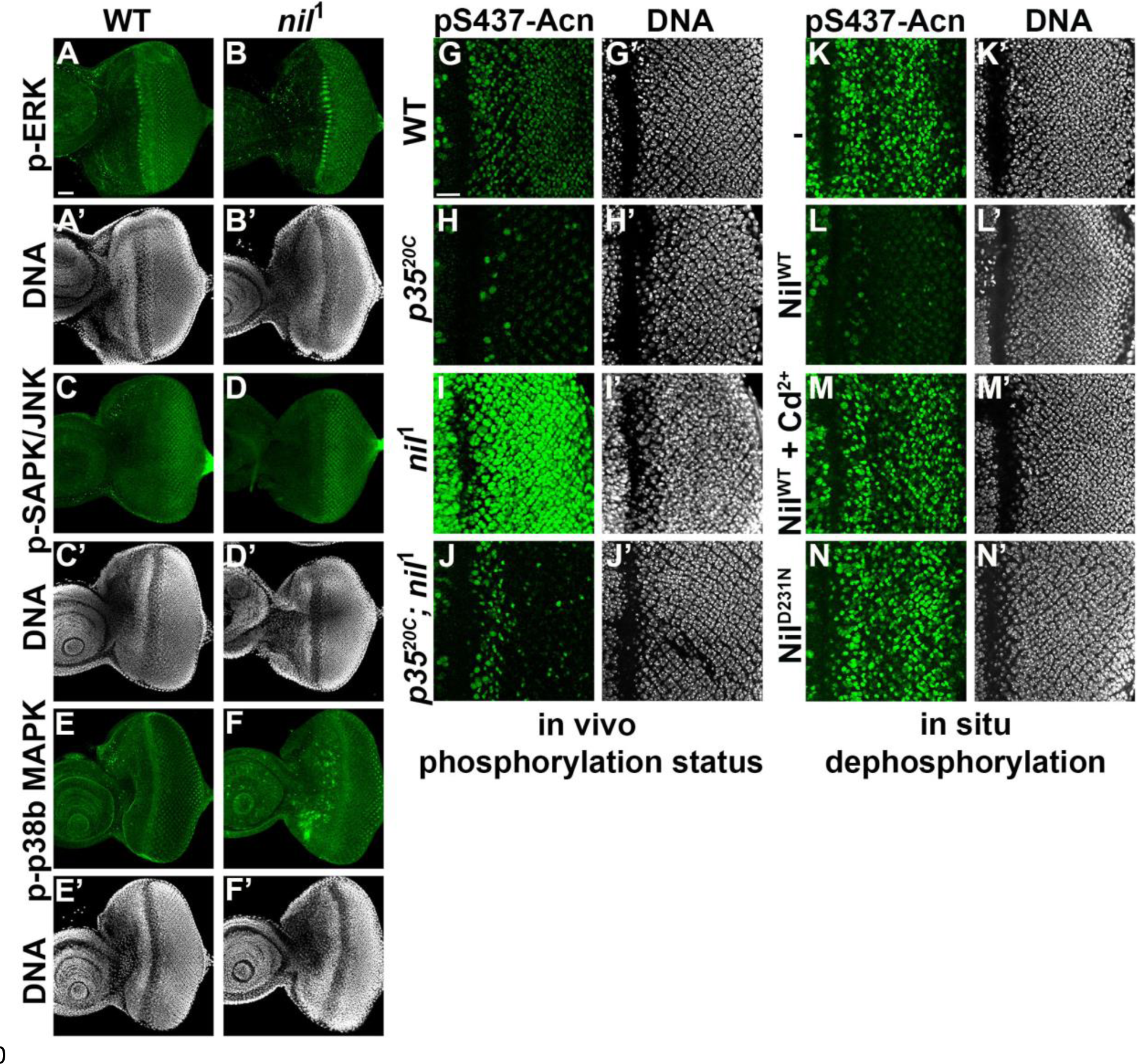
Nil dephosphorylates Acinus counteracting Cdk5-p35-mediated phosphorylation. (A–F) Projection of confocal micrographs of *w*^1118^ and *nil*^1^ larval eye discs stained for p-ERK (green) and DNA (A- B’), p-SAPK/JNK (green) and DNA (C-D’), p-p38b MAPK (green) and DNA (E-F’). (G-J) Projection of confocal micrographs of larval eye discs stained for pS437-Acn (green) and DNA from *w*^1118^ (G, G’), *p35*^20C^ (H, H’), *nil*^1^ (I, I’), *p35*^20C^*; nil^1^* (J, J’). (K-N) Projection of confocal micrographs of eye discs from Acn^WT^ larvae stained for pS437-Acn (green) and DNA without (K,K’) or after in-situ dephosphorylation with wild-type Nil (L,L’), wild-type Nil + 100 µM CdCl2 (M,M’), or inactive Nil^D231N^ (N,N’). Scale bar in A is 40 µm for A-F. Scale bar in G is 20 µm for G-N. Genotypes are listed in Supplemental Table 3.

By contrast, the Cdk5-p35 kinase complex can directly phosphorylate Acn-S437 (Nandi et al., 2017). To further test whether kinases other than Cdk5-p35 contribute to elevated pS437-Acn in *nil*^1^ animals, we examined Acn S437 phosphorylation in *nil*^1^; *p35*^20C^ double mutants. With the exception of dividing cells close to the morphogenetic furrow, *nil*^1^; *p35*^20C^ double mutant eye discs failed to display the pS437-Acn levels (Figure 4J) observed in *nil*^1^ eye discs (Figure 4I) and instead were similar to *p35*^20C^ mutants (Figure 4H,J). Taken together, these data indicate that the Nil phosphatase counteracts Acn-phosphorylation by the Cdk5-p35 kinase complex rather than inactivating a stress-responsive MAPK.

### Nil phosphatase contributes to Cd^2+^ toxicity and neurodegenerative stress

Cd^2+^ targets the M1 metal ion binding site of mammalian PPM1 phosphatases and efficiently inhibits them (Pan et al., 2013). To test whether the Nil phosphatase is inhibited by Cd^2+^ as well, we developed an in-situ assay. Eye-discs from Acn^WT^ larvae were fixed and detergent-treated to preserve the phosphorylation status of Acinus which was detected by pS437-Acn staining (Figure 4K). In the fixed tissue, pS437-Acn was dephosphorylated by purified Nil phosphatase (Figure 4L), but not by Cd^2+^-inhibited Nil (Figure 4M) or inactive Nil^D231N^ (Figure 4N), consistent with sensitivity to Cd^2+^ inhibition being conserved in the Nil phosphatase.

Cd^2+^-induced cytotoxicity is associated with oxidative stress (Branca et al., 2020), which can be reduced by elevated levels of basal autophagy (Galati et al., 2019; Yun et al., 2020). To test a possible role of the Nil phosphatase in Cd^2+^-induced cellular stress responses, we examined whether pS437-Acn levels increase upon exposure to environmental Cd^2+^. We found that eye discs from wild-type larvae grown in 100 µM Cd^2+^ displayed elevated phosphorylation of Acn at S437 with a concomitant increase in the number of ATG8a positive punctae (Figure 5A-D,I).

**Figure 5.**
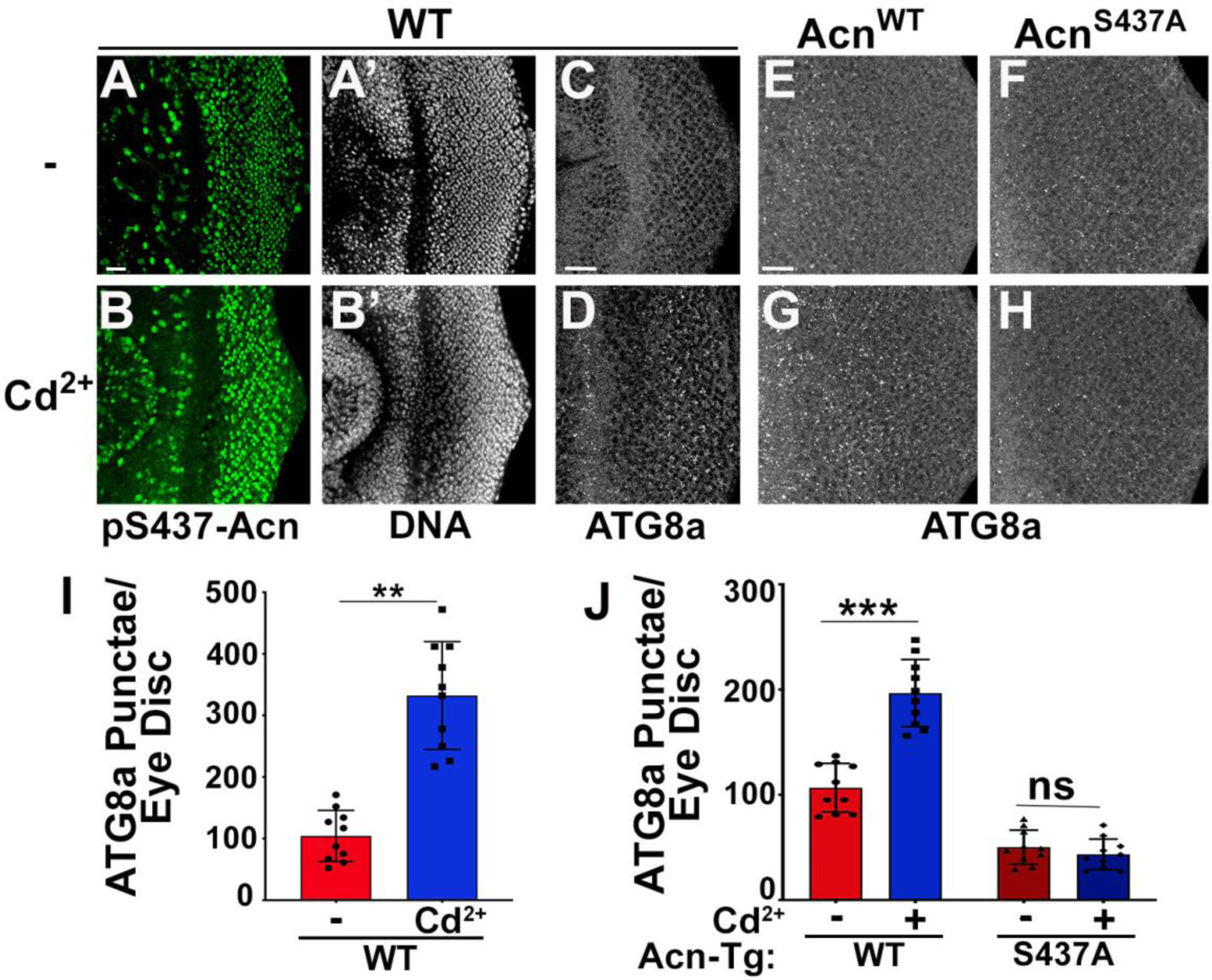
Acinus-S437 phosphorylation is required for Cd^2+^-induced autophagy. (A-B) Projection of confocal micrographs of *w*^1118^ larval eye discs stained for pS437-Acn (green) and DNA from either 100 µM CdCl2 treated (B) or untreated (A) larvae. (C-D) Projection of confocal micrographs of fed *w*^1118^ larval eye discs either 100 µM CdCl2 treated (D) or untreated (C) stained for Atg8a. (E-H) Projection of confocal micrographs of fed Acn^WT^ and Acn^S437A^ larval eye discs either 100 µM CdCl2 treated (G, H) or untreated (E, F) stained for Atg8a. (I) Quantification of Atg8a punctae in either 100 µM CdCl2 treated or untreated *w*^1118^ larval eye discs. Data are from 10 larvae from three experimental repeats. Bar graphs show mean ±SD. **p<0.01 (J) Quantification of Atg8a punctae in either 100 µM CdCl2 treated or untreated Acn^WT^ and Acn^S437A^ larval eye discs. Data are from 10 larvae from three experimental repeats. Bar graphs show mean ±SD. ns, not significant; ***p<0.001. Scale bar in A is 20 µm for A-H. Genotypes are listed in Supplemental Table 3.

Cd^2+^ may also effect other signaling pathways with the potential to alter autophagy. We therefore wanted to test whether elevated Acn-S437 phosphorylation is necessary for Cd^2+^- induced autophagy. For this purpose, we analyzed the effect of Cd^2+^ on basal autophagy in larvae expressing either Acn^WT^ or Acn^S437A^ under control of the Acn promoter in an *acn* null background (Nandi et al., 2017). We observed an increase in ATG8a punctae in fed eye discs from Acn^WT^ larvae grown in 100 µM Cd^2+^ similar to wild-type animals (Figure 5E,G,J). By contrast, Cd^2+^ exposure failed to elevate basal autophagy in the phospho-inert Acn^S437A^ mutants (Figure 5F,H,J) indicating that Acn-S437 phosphorylation is necessary for an autophagic response to Cd^2+^ exposure. Taken together, these data suggest exposure to Cd^2+^ can elevate pS437-Acn levels and enhance basal autophagy flux by deactivating Nil phosphatase.

These findings motivated us to further investigate a possible role of the Nil phosphatase in Cd^2+^-induced cytotoxicity. We exposed wild-type and *nil*^1^ flies to varying concentrations of Cd^2+^ and compared their survival. Compared to wild type, median survival time for *nil*^1^ mutants increased by 2.5 days on exposure to 250 µM Cd^2+^, and 5 days at 375 µM Cd^2+^ (Figure 6C,D). Interestingly, this difference in susceptibility to Cd^2+^ poisoning was confined to a narrow concentrations range: in the absence of Cd^2+^ (Figure 6A, Figure 6-figure supplement 1), or at higher concentrations (500 µM, Figure 6E), wild type and *nil*^1^ mutants were not different in their survival. These data suggest that the elevated autophagy in *nil*^1^ mutants helps the animals to cope with low levels of Cd^2+^-induced oxidative stress, but is overwhelmed at higher levels.

**Figure 6.**
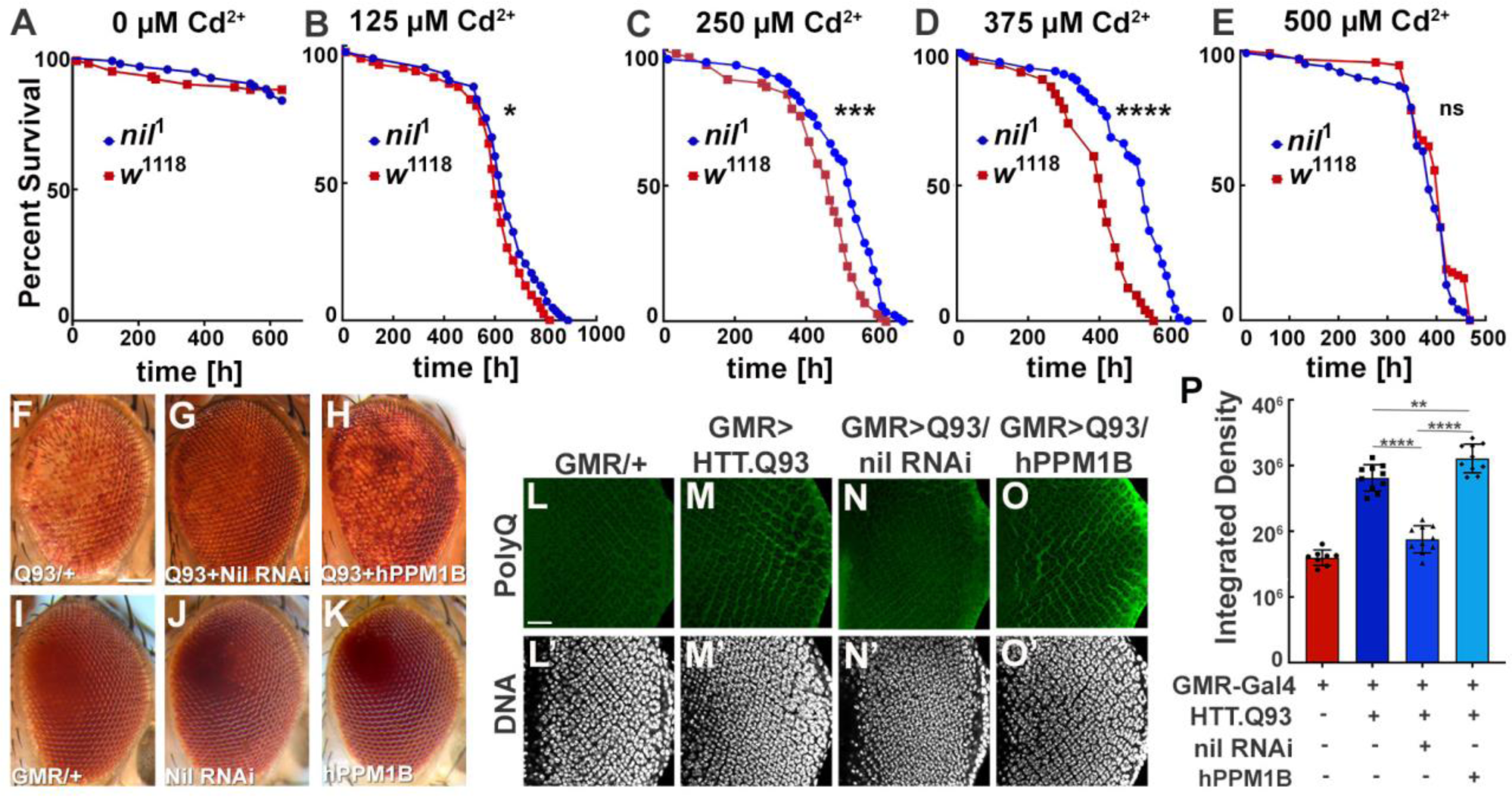
Loss of Nil function provides partial protection against Cd^2+^ poisoning and proteostasis stress. (A-E) Survival curves for *w*^1118^ and *nil*^1^ adult male flies either untreated (A) or treated with 125 µM CdCl2 (B), 250 µM CdCl2 (C), 375 µM CdCl2 (D), 500 µM CdCl2 (E). Log-rank comparisons revealed significant differences between survival curves: ns, not significant; *p<0.05; ***p<0.001; ****p<0.0001 (F–K) Micrographs of adult eyes in which GMR-Gal4 drives expression of UAS-HTT^ex1^.Q93 (F), UAS-HTT^ex1^.Q93 + UAS-Nil RNAi (G), UAS-HTT^ex1^.Q93 + UAS-hPPM1B (H), UAS-Nil RNAi (J), UAS-hPPM1B (K), and (I) represents GMR-Gal4 control. Scale bar in F is 100 µm for F-K. (L-O) Projection of confocal micrographs of larval eye discs stained for PolyQ (green) and DNA from GMR-Gal4 (L, L’), GMR-Gal4, UAS-HTT^ex1^.Q93 (M, M’), GMR-Gal4, UAS-HTT^ex1^.Q93 + UAS-Nil RNAi (N, N’), GMR-Gal4, UAS-HTT^ex1^.Q93 + UAS-hPPM1B (O, O’). (P) Quantification of PolyQ accumulation in eye discs of the indicated genotypes from a constant area starting 2–3 rows posterior to the furrow containing around 100 ommatidial clusters. Bar graphs show mean ± SD of integrated densities from 10 larvae taken out of three experimental repeats. **p<0.01; ****p<0.0001. Scale bar in L is 20 µm for L-O. Genotypes are listed in Supplemental Table 3.

Cdk5-p35-mediated AcnS437 phosphorylation alleviates proteostatic stress in Drosophila models of neurodegenerative diseases (Nandi et al., 2017). Therefore, we wondered whether loss of Nil phosphatase function may reduce neurodegenerative stress. Eye-specific expression of Huntingtin-polyQ polypeptides (HTT.Q93) results in neuronal degeneration reflected by depigmentation of the adult eye (Figure 6F,I, Supplemental Table 2) as previously shown (Xu et al., 2015). This depigmentation phenotype is suppressed by knockdown of the Nil phosphatase (Figure 6G,J, Supplemental Table 2) but only marginally altered by overexpression of PPM1B, the human homolog of Nil, (Figure 6H,K, Supplemental Table 2). To more directly asses the effect of Nil phosphatase on the accumulation of polyQ proteins, we stained wandering larval eye discs for polyQ proteins. GMR-driven expression of HTT.Q93 resulted in accumulation of polyQ in eye discs a few rows posterior to the furrow (Figure 6L,M,P). Knocking down Nil phosphatase expression significantly reduced polyQ accumulation posterior to the furrow (Figure 6N,P). By contrast, overexpression of the human PPM1B phosphatase further enhanced the polyQ load (Figure 6I,P). This is consistent with the data above that show elevated autophagy in *nil*^1^ eye discs in combination with the known role of autophagy in the clearance of protein aggregates (Menzies et al., 2017). Taken together, these data suggest that PPM1-type phosphatases play an important role in regulating cellular responses to Cd^2+^ toxicity and neurodegenerative stress.

## Discussion

We previously identified Acn as a signaling hub that integrates multiple stress response pathways to regulate autophagy (Nandi and Krämer, 2018). Starvation-independent autophagy is elevated in response to Acn being stabilized either by inhibition of its Caspase- 3 mediated cleavage (Nandi et al., 2014) or by Cdk5-p35-mediated phosphorylation (Nandi et al., 2017). Here, we extend this concept to Nil-regulated dephosphorylation of Acn. We show that among the serine/threonine phosphatases encoded in the Drosophila genome the PPM-type phosphatase Nil is specifically responsible for counteracting Acn phosphorylation by the Cdk5-p35 complex, a function conserved in the human PPM1B homolog of Nil. We used three different methods to interfere with Nil function: RNAi-induced knockdown, the CRISPR/Cas9-generated *nil*^1^ null allele, or Cd^2+^-mediated inhibition of Nil. All three yielded increased pS437-Acn levels and elevated autophagic flux. The increased autophagy, in agreement with its well-established beneficial functions in other contexts (Levine and Kroemer, 2019), extended survival time for flies exposed to Cd^2+^-laced food and reduced polyQ load in a Drosophila model of Huntington’s disease. Thus, regulation of PPM-type phosphatase function may play an underappreciated role in the regulation of quality-control autophagy.

The family of PPM-type phosphatases is represented in the genomes of eukaryotes from yeast to humans and individual family members are highly conserved across phyla (Kamada et al., 2020). Despite roles of these phosphatases in diverse physiological contexts, including the regulation of metabolism, cell cycle progression, immunological responses and other stress responses, the in-vivo exploration of their functions in animal models lacks behind that of their kinase counterparts. For example, the *Drosophila melanogaster* genome encodes 15 isoforms of PPM-type phosphatases (Kamada et al., 2020), but to our knowledge only two of them have previously been characterized in detail using null alleles. Interestingly, these studies revealed specific regulatory roles for both phosphatases: (i) Pdp (encoded by the *pyruvate dehydrogenase phosphatase* gene) dephosphorylates the Mad signal transducer and thereby negatively regulates BMP/Dpp signaling (Chen et al., 2006), (ii) the Alphabet phosphatase, similar to its human PPM1A/B homologs, regulates responses to developmental, oxidative or genotoxic stresses through dephosphorylation of different MAP Kinases (Baril et al., 2009; Baril and Therrien, 2006). Alphabet, among Drosophila phosphatases, is the one most similar to Nil (Kamada et al., 2020). Therefore, we tested whether Nil also affects phosphorylation of stress-activated MAP kinases and thereby indirectly alters Acn phosphorylation. We could not find evidence supporting this possibility. In *nil*^1^ mutants, pS437-Acn phosphorylation still depended on Cdk5-p35 activity and levels of phosphorylated forms of ERK, Jun or p38 kinases appeared unchanged. Together, these findings argue against a contribution of stress-activated kinases, other than Cdk5-p35, in Nil’s effect on regulating phospho-Acn levels and autophagy.

Interestingly, other PPM-type phosphatases have also been implicated in the regulation of autophagy. In yeast, the Ptc2 and Ptc3 phosphatases redundantly regulate autophagy through the dephosphorylation of Atg1 and its binding partner Atg13 (Memisoglu et al., 2019). In mammalian cells, genotoxic stress activates PPM1D to dephosphorylate ULK1 and activate autophagy (Torii et al., 2016). In both cases, phosphatase activity counteracts the mTOR-mediated inhibition of autophagy. We do not know whether the Nil phosphatase affects other autophagy-related proteins in addition to Acn but, at least in the context of cadmium-induced autophagy, a critical step is the inhibition of Acn-S437 dephosphorylation by Nil, as cadmium can no longer induce autophagy in the phospho-inert *acn*^S437A^ background.

Cadmium is an environmental pollutant which, unlike many other metal ions, has no known biological role. Toxic effects of elevated cadmium levels can manifest as kidney or skeletal diseases and have been linked to multiple cancers (Templeton and Liu, 2010; WHO, 2020) and neurodegenerative disorders (Branca et al., 2020). The effects of cadmium on autophagy appear to be complex. Our data, in agreement with previous studies (So et al., 2018; Zhang et al., 2016), show elevated cadmium to induce autophagy with resulting cytoprotective effects, and we identify the inhibition of Nil-mediated Acn dephosphorylation as a key mechanism for this induction. In other contexts, especially cancer cells, cadmium appears to inhibit autophagy (Liang et al., 2021), suggesting that Cadmium interacts with a distinct signaling network in those cells. Interestingly, regulation of autophagy by Acn is independent of mTor signaling. The Acn^S437A^ mutation interferes with autophagy induction by Cadmium or proteostasis stress, but does not block the mTor-dependent activation of autophagy upon amino-acid starvation (Nandi et al., 2017).

While there is now ample support for a role of Acn in regulating autophagy (Haberman et al., 2010; Nandi and Krämer, 2018; Nandi et al., 2017; Orvedahl et al., 2011), likely upstream of Atg1/ULK kinases (Tyra et al., 2020), the specific mechanism is not clear yet. Nil’s localization to the nucleus is consistent with effects on the well-established role of Drosophila and mammalian Acn proteins in alternative splicing (Deka and Singh, 2019; Hayashi et al., 2014; Michelle et al., 2012; Murachelli et al., 2012; Nandi and Krämer, 2018; Rodor et al., 2016; Schwerk et al., 2003; Singh et al., 2010). Alternatively, the subset of Nil protein localizing close to endolysosomal compartments and autophagosomes points to an alternative possibility of a more direct role of phosphorylated Acn in regulating autophagic flux. Future work will be aimed at distinguishing between these possibilities.

## Author Contributions

NN conceived and executed experiments, analyzed data and drafted an early manuscript. ZZ executed experiments and analyzed data. CT executed experiments, HK conceived and executed experiments and analyzed data. All co-authors participated in revising the manuscript.

## Acknowledgements

We thank members of the Krämer lab for helpful comments to the manuscript and technical assistance. We thank Drs. Edward Giniger, National Institute of Neurological Disorders and Stroke, Bethesda, Maryland, Robin Hiesinger, Free University Berlin, Berlin, Germany, the Vienna Drosophila Resource Center, and the Bloomington Drosophila Stock Center (NIH P40OD018537) for flies, and Akira Nakamura, RIKEN Center for Developmental Biology, Kobe, Japan, the Developmental Studies Hybridoma Bank at The University of Iowa for antibodies. This work was funded by NIH grants R01EY010199 and R21EY030785.

## Methods

### CONTACT FOR REAGENT AND RESOURCE SHARING

Further requests for information or resources and reagents should be directed to and will be fulfilled by the Lead Contact, Helmut Kramer

### EXPERIMENTAL MODEL

Fly stocks were maintained at room temperature under standard conditions. Bloomington *Drosophila* Stock Center provided Da-Gal4, GMR-Gal4 driver lines, *w*^1118^, serine-threonine phosphatase RNAi lines and UAS-hPPM1B (BS76916). Other fly strains used were *p35^20C^*, which deletes ∼90% of the *p35* coding region including all sequences required for binding to and activating Cdk5 (Connell-Crowley et al., 2007), a kind gift from Edward Giniger, National Institute of Neurological Disorders and Stroke, Bethesda, Maryland, YFP^MYC^-Rab5 (Dunst et al., 2015) and UAS-Htt-exon1-Q93 (Steffan et al., 2001), abbreviated UAS-Htt.Q93, was a gift from Robin Hiesinger, Free University Berlin, Berlin, Germany.

*nil*^1^ null mutants and the endogenously tagged *nil* gene were generated essentially as described (Stenesen et al., 2015) using tools available from the O’Connor-Giles, Wildonger, and Harrison laboratories (Gratz et al., 2013). Specifically, gRNAs (see Supplemental Table 4) were introduced into the pU6-BbsI vector and co-injected with the appropriate template plasmid for homologous repair. Embryo injections were done by Rainbow Transgenic Flies (Camarillo, CA), and the resulting potentially chimeric adult flies were crossed with *w*^1118^ flies.

For the *nil*^1^ null allele, the template plasmid was assembled in the pHD-DsRed backbone using approximately 1kb PCR-amplified 5’ and 3’ homology arms. Their progeny, potential founders, were crossed to balancer stocks and resulting flies with eye-specific DsRed expression (Bier et al., 2018) were selected, balanced and homozygotes collected for further analysis.

For the *nil*^Ty1_Gal4^ allele, the template plasmid was assembled in pBS-3xTy1-T2A- Gal4. Flanking homology 5’ and 3’ arms of approximately 1kb were synthesized as gBlocks with mutations in the gRNA target sites. Progeny from the initial cross with *w*^1118^ flies, potential founders, were crossed to flies containing a 20xUAS-6xmCherry-HA cassette, Bloomington stock: 52267, (Shearin et al., 2014). Males with elevated abdominal mCherry expression were identified and balanced. Both alleles, *nil*^1^ and *nil*^Ty1_Gal4^, were confirmed by sequencing of PCR products generated with one primer outside the homology arms.

Transgenic flies were generated by BestGene, Inc. DNA constructs related to genomic *acn* were generated by standard mutagenesis of a 4-kb Acn DNA fragment sufficient for genomic rescue (Haberman et al., 2010), confirmed by sequencing, cloned into an Attb vector, and inserted into the 96F3 AttP landing site (Venken et al., 2006). UAS-Acn transgenes inserted into 43A1 landing site were previously described (Nandi et al., 2017).

To maximize knockdown efficiency experiments with UAS-RNAi transgenes were performed at 28°C.

Life span were analyzed as described previously (Nandi et al., 2014). Briefly, to measure life spans, males that emerged within a 2-day period were pooled and aged further for an additional 3 days, and then placed in demographic cages and their survival at 25°C was recorded every day. Around 100 flies were kept in each demographic cage with three replicates for each genotype. Food vials were changed every other day, and dead flies were counted and removed.

For Cd^2+^ exposure of Drosophila larvae, fly eggs were allowed to develop on an apple juice agar plate containing the desired concentration of CdCl2 at 25°C. For analyzing Cd^2+^ toxicity, adult flies are kept in standard fly food with the desired concentrations of CdCl2 at 25°C. Males that emerged within a 2-day period were collected and aged for 3 more days, before being placed in demographic cages to record their survival every day. Around 50 flies are kept in each demographic cage with three replicates for each set. Food vials with the desired concentrations of CdCl2 were changed every other day, and dead flies were counted and removed.

#### Histology

Eye micrographs were obtained at 72X magnification on a SteREO-microscope (SteREO Discovery.V12; Carl Zeiss) with a camera (AxioCam MRc 5; Carl Zeiss) using AxioVision image acquisition software (Carl Zeiss). Images of fly eyes are a composite of pictures taken at multiple z positions and compressed using CZFocus (Carl Zeiss) or Helicon Focus (Helicon Soft) software.

#### Biochemistry

Quantitative RT-PCR was used to measure knockdown efficiencies as previously described (Akbar et al., 2011). In short, RNA was isolated using TRIZOL (Ambion) according to the manufacturer’s protocol. High-Capacity cDNA Reverse Transcription kit (Applied Biosystems) was then used to reverse transcribe 2 µg RNA with random hexamer primers. Quantitative PCR was performed using the Fast SYBR Green Master Mix in a real-time PCR system (Fast 7500; Applied Biosystems). Each data point was repeated three times and normalized for the data for ribosomal protein 49 (RP49). Primers are listed in Supplemental Table 4.

For immunoblot experiments, 25 adult fly heads were homogenized in 250 µl lysis buffer (10% SDS, 6 M urea, and 50 mM Tris-HCl, pH 6.8) at 95°C, boiled for 2 min, and spun for 10 min at 20,000x*g*. 10 µl lysate from larvae were separated by SDS-PAGE, transferred to nitrocellulose membranes, blocked in 3% non-fat dried milk and probed with rabbit anti- ATG8a (1:1000), rabbit anti-hook (1:5000), mouse anti-actin (JLA20) (1:2000) and mouse anti-Ty1 clone BB2 (1:2000). Bound antibodies were detected and quantified using IR-dye labelled secondary antibodies (1:15,000) and the Odyssey scanner (LI-COR Biosciences).

Pre-stained molecular weight markers (HX Stable) were obtained from UBP-Bio.

#### Immunofluorescence

Whole-mount tissues for immunofluorescence staining were set up as described previously (Nandi et al., 2017). Briefly, dissected tissues after fixation in periodate-lysine-paraformaldehyde were washed in 1X PBS, permeabilized with 0.3% saponin in 1X PBS (PBSS), blocked with 5% goat serum in PBSS, and stained with the specified primary antibodies: rabbit anti-pS437Acn (1:1000, Nandi et al., 2017), guinea pig anti-Acn (1:1000, Nandi et al., 2014), mouse anti-V5 (1:500, Invitrogen), mouse anti-Ty1 clone BB2 (1:500, Invitrogen), rabbit anti-Arl8 at 1:300, (Boda et al., 2019), rabbit anti-Rab7 (1:3000, Tanaka et al., 2008, a kind gift from Akira Nakamura, RIKEN Center for Developmental Biology, Kobe, Japan), rabbit anti-phospho-p44/42 MAPK (Erk1/2) (Thr202/Tyr204) (D13.14.4E) (1:200, CST), mouse anti-phospho-SAPK/JNK (Thr183/Tyr185) (G9) (1:200, CST), rabbit anti-phospho-p38 MAPK (Thr180/Tyr182) (D3F9) (1:500, CST), mouse-anti 1C2 (1:1,000; MAB1574; EMD Millipore), rabbit anti-GABARAP (1:200; Abcam, ab109364), which detects endogenous Atg8a (Kim et al., 2015). Next, the tissues stained with primary antibodies overnight were washed and stained with secondary antibodies conjugated to Alexa Fluor 488, 568, or 647 (1:500; Molecular Probes) and mounted in Vectashield containing DAPI (Vector Laboratories). Fluorescence images were captured with 63X, NA 1.4 or 40X, NA1.3 or 20X, NA0.8 Plan Apochromat lenses on an inverted confocal microscope (LSM 710; Carl Zeiss). Confocal Z-stacks were acquired at 1-µm step size.

For analysing autophagy flux 72-hours old larvae were transferred to fresh medium containing 3 mg/ml chloroquine (Sigma) as described previously (Low et al., 2013).

Z-projections of three optical sections for fat body tissue and eight optical sections for eye discs and antennal discs, each 1 µm apart were used to quantify Atg8a punctae using Imaris software (Bitplane). For fat bodies, the number of punctate quantified represent per fat body cell. Integrated densities for polyQ in identical areas posterior to the morphogenetic furrow of eye discs were quantified using Image J software.

Digital images for display were bring into Photoshop (Adobe) and tuned for gain, contrast, and gamma settings.

All immunofluorescence experiments were repeated at least three times with at least three samples each.

#### In-situ dephosphorylation assay

Puromycin-selectable plasmids for the expression of C-terminally Twinstreptag-Flag- tagged Nil^WT^ and Nil^D231N^ proteins under control of the metallothionine promoter were transfected in S2 cells. Selected pools of 6x10^6^ cells were induced with 0.7 mM CuSO4 for 16 hours. Cells were lysed in RIPA buffer, Nil proteins bound to MagStrep beads and washed following the manufacturer’s instructions (IBA GmbH, Göttingen, Germany). Nil proteins were eluted using 50 mM Biotin in elution buffer (50 mM Tris pH8.5, 40 mM NaCl, 0.1 mM EGTA, 1 mM DTT).

The in situ dephosphorylation assay was slightly modified from the method described (Nandi et al., 2017). Briefly, dissected third instar larval carcasses were fixed) in periodate-lysine-paraformaldehyde, permeabilized using PBSS and treated with either Nil^WT^ or Nil^WT^ + 100 µM CdCl2 or the inactive phosphatase Nil^D231N^ in 1X phosphatase assay buffer (40 mM MgCl2, 40 mM MnCl2, 50 mM Tris pH 8.5, 40 mM NaCl, 0.1 mM EGTA, 1X EDTA free Protease inhibitor (Roche) and 1 mM PMSF) for 3 h at 37°C. Following the phosphatase reaction, eye discs were washed 3x in PBSS and stained for pS437-Acn, mounted and imaged as described above.

#### Statistical methods

Statistical significance was determined in Prism using one-way analysis of variance for multiple comparisons, followed by Tukey’s test and log-rank for survival assays. We used two-way analysis of variance for multiple comparisons, followed by Bonferroni’s test for individual comparisons to separate effects of treatment and genetic background. Bar graphs generated from this analysis demonstrate means ±SD. For quantifications of fluorescence images at least three independent experiments were used. P values smaller than 0.05 are considered significant, and values are indicated with one (<0.05), two (<0.01), three (<0.001), or four (<0.0001) asterisks.

## Supplemental Information

**Figure 1-figure supplement 1.**
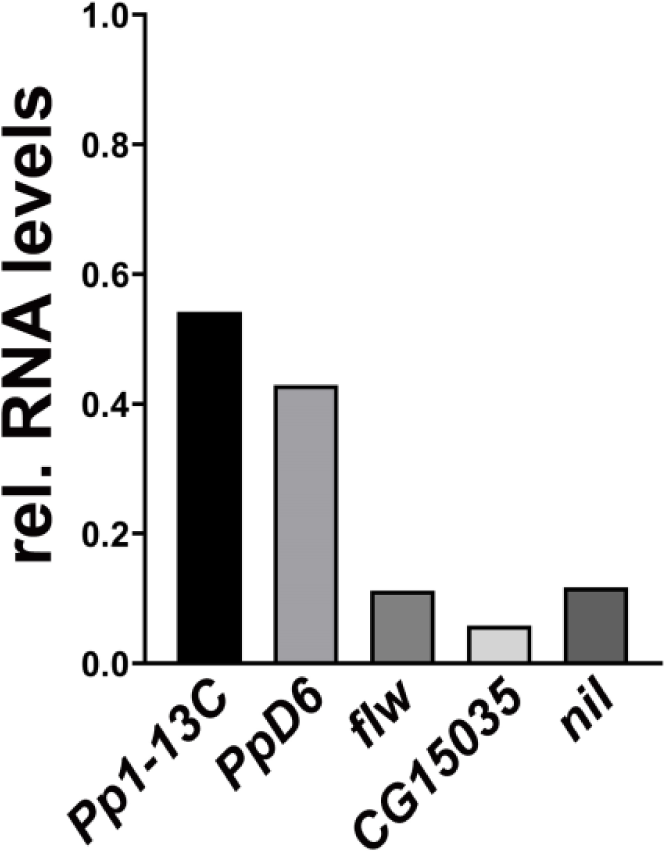
qPCR analysis of phosphatase genes interacting with Acinus. Da-Gal4 driven knockdown of indicated phosphatases in wandering third instar larvae and analysis of the fold decrease in transcript level by quantitative RT-PCR. The data is normalized to levels of RP49 transcription. Genotypes are listed in Supplemental Table 3.

**Figure 2-figure supplement 1.**
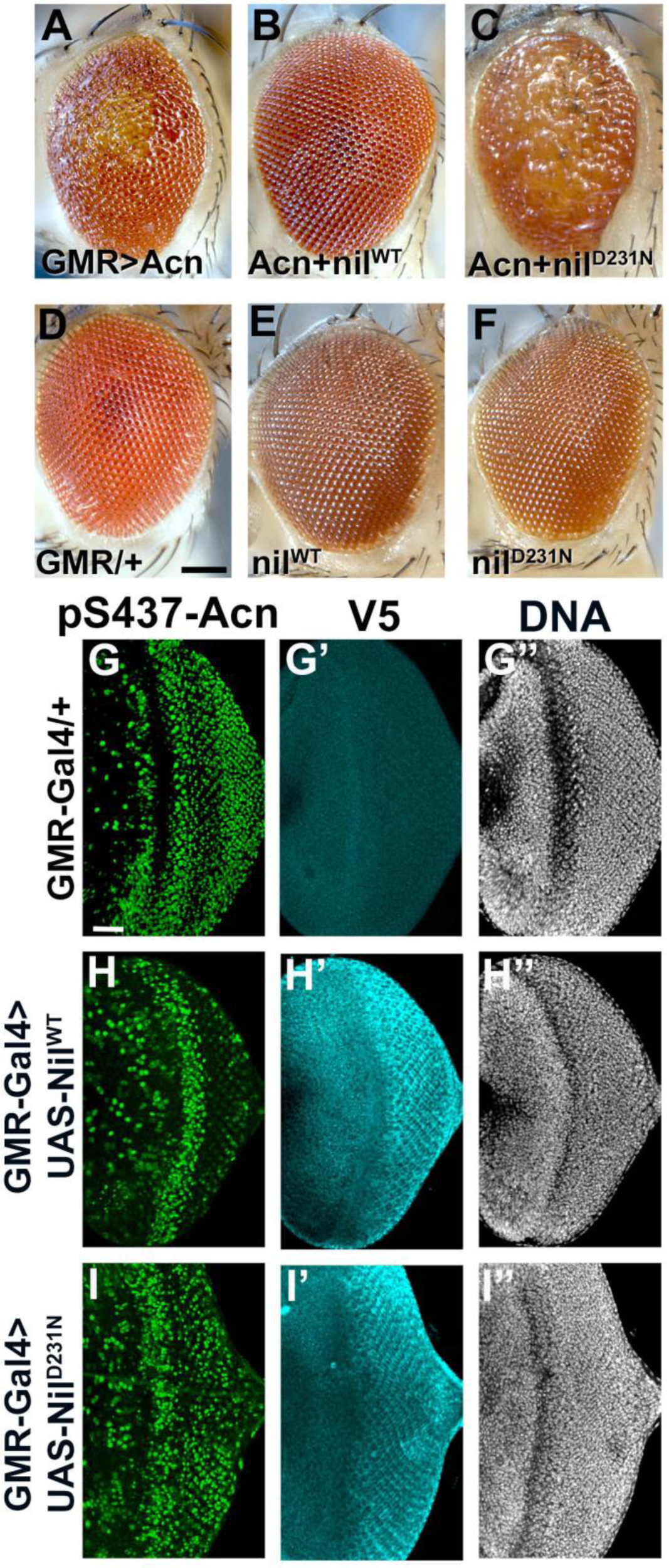
Effects of Nil overexpression depend on its phosphatase activity. (A–F) Micrographs of eyes in which GMR-Gal4 drives expression of Acn^WT^ (A), Acn^WT^ + Nil^WT^ (B), Acn^WT^ + Nil^D231N^ (C), Nil^WT^ (E), Nil^D231N^ (F) and K represents GMR-Gal4 control. Scale bar in D is 100 µm for A-F. (G–I) Projection of confocal micrographs of larval eye discs stained for pS437-Acn (green), V5 (cyan) and DNA from GMR-Gal4 (G-G’’), GMR-Gal4, UAS- Nil^WT^ (H-H’’), GMR-Gal4, UAS- Nil^D231N^ (I-I’’). Nil transgenes are V5 tagged. Scale bar in G is 20 µm for G-I. Genotypes are listed in Supplemental Table 3

**Figure 2-figure supplement 2.**
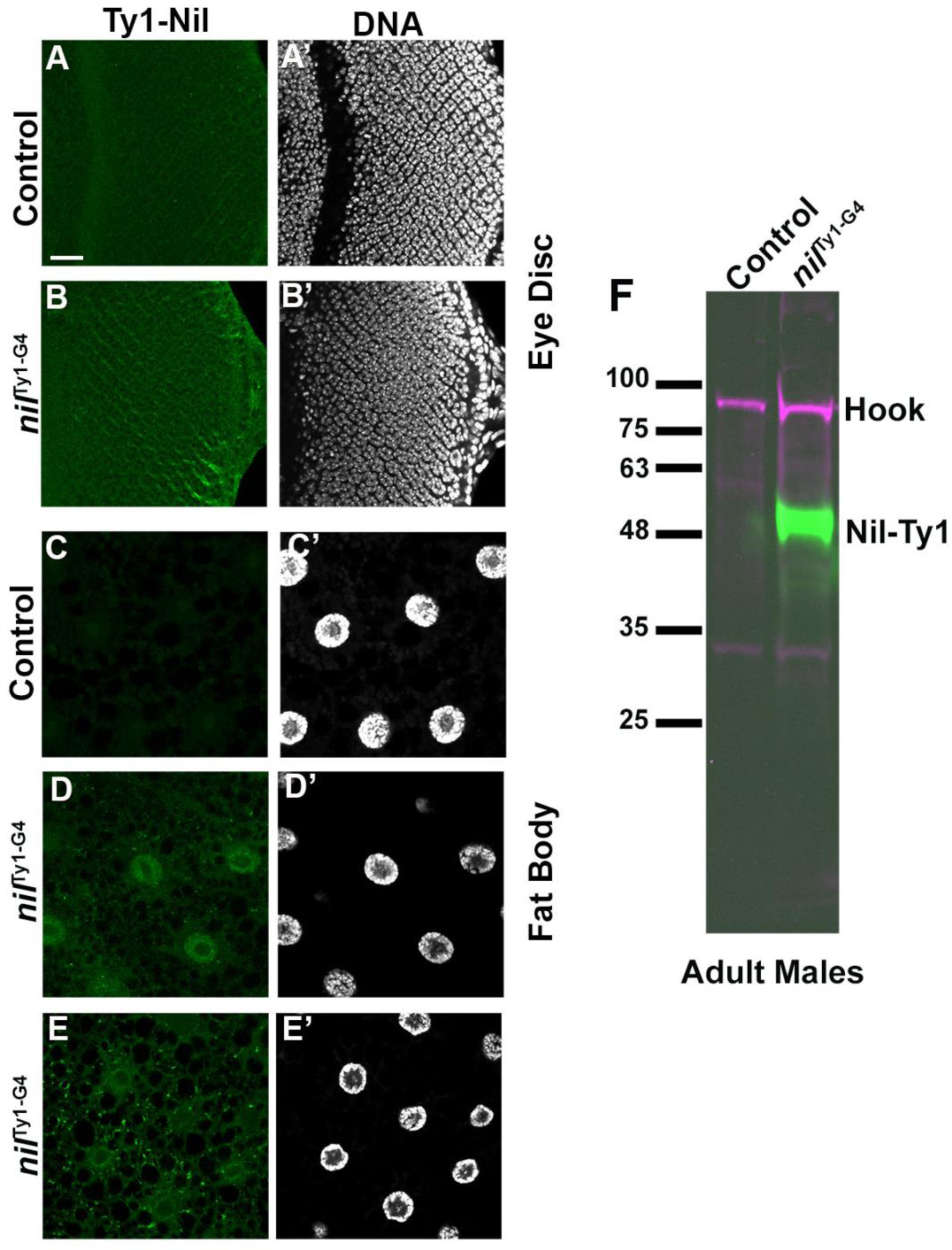
Detection of Ty1-tagged endogenous Nil protein. (A-E) Projection of confocal micrographs of larval eye discs (B) and fat bodies (D, E) from nil^Ty1-G4^ stained for Ty1(green). No Ty1 (green) staining is observed in appropriate controls (A, C). (F) Western blot of lysates from nil^Ty1-G4^ and appropriate control adult male flies probed for Ty1 and Hook as control. Scale bar in A is 20 µm for A-E. Genotypes are listed in Supplemental Table 3.

**Figure 6-figure supplement 1 .**
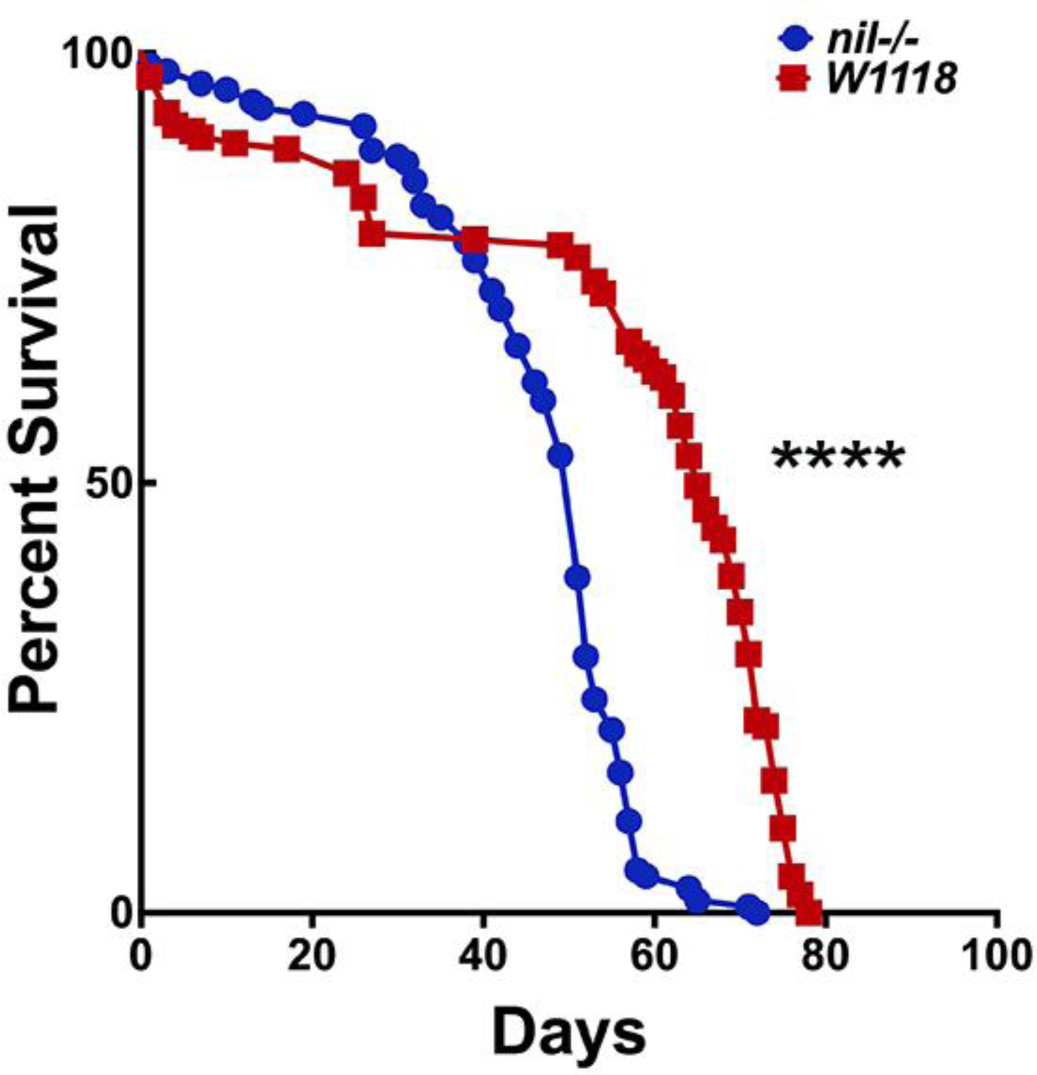
Survival curves for adult male *w*^1118^ and *nil^1^* flies. Log-rank comparisons revealed significant difference between survival curves: ****p<0.0001. Genotypes are listed in Supplemental Table 3.

**Supplemental Table I.** Genetic interactions of Acinus gain-of-function with different phosphatases

| GMR-Gal4 driven transgenes | Normal | Mild Errors | Rough Eyes | Severely Rough | Avg. Score | n = flies scored | Enhancement or Suppression |
| --- | --- | --- | --- | --- | --- | --- | --- |
| Acinus (Acn) | 6 | 5 | 3 | 12 | 2.8 | 26 | N/A |
| GMR-Gal4 | 56 | 10 | 0 | 0 | 1.2 | 66 | N/A |
| Acn + Alphabet (31398) | 12 | 17 | 12 | 26 | 2.8 | 67 | 0% |
| Alphabet (31398) | 31 | 21 | 0 | 0 | 1.4 | 52 |  |
| Acn + Alphabet (40873) | 1 | 6 | 30 | 53 | 3.2 | 90 | 33.24% |
| Alphabet (40873) | 38 | 33 | 20 | 3 | 1.9 | 96 |  |
| Acn+ CG6036 (65115) | 3 | 4 | 3 | 27 | 3.5 | 37 | 58.77% |
| CG6036 (65115) | 42 | 5 | 0 | 0 | 1.1 | 47 |  |
| Acn + CG6036 (105568) | 3 | 23 | 5 | 24 | 2.9 | 55 | 5.56% |
| CG6036 (105568) | 15 | 9 | 12 | 18 | 1.8 | 54 |  |
| Acn + CG7115 (60015) | 8 | 10 | 8 | 15 | 2.7 | 41 | -5.39% |
| CG7115 (60015) | 47 | 20 | 0 | 0 | 1.3 | 67 |  |
| Acn + CG7115 (103354) | 5 | 22 | 2 | 17 | 3.0 | 46 | 11.11% |
| CG7115 (103354) | 23 | 23 | 10 | 0 | 1.8 | 56 |  |
| Acn + CG10376 (41907) | 15 | 13 | 12 | 5 | 2.2 | 45 | -58.56% |
| CG10376 (41907) | 64 | 8 | 0 | 0 | 1.1 | 72 |  |
| Acn + CG10376 (57036) | 3 | 6 | 0 | 26 | 3.4 | 35 | 25% |
| CG10376 (57036) | 16 | 7 | 12 | 0 | 2.4 | 31 |  |
| Acn + CG10417 (39051) | 16 | 27 | 3 | 29 | 2.3 | 75 | -41.96% |
| CG10417 (39051) | 90 | 13 | 0 | 0 | 1.1 | 103 |  |
| Acn + CG12091 (40936) | 23 | 21 | 42 | 0 | 2.7 | 86 | 7.83% |
| CG12091 (40936) | 74 | 13 | 0 | 0 | 1.2 | 87 |  |
| Acn + CG12091 (105722) | 5 | 8 | 0 | 20 | 3.1 | 33 | 17.34% |
| CG12091 (105722) | 42 | 25 | 5 | 0 | 1.7 | 70 |  |
| Acn + CG15035 (60404) | 1 | 5 | 0 | 28 | 3.6 | 34 | 74.95% |
| CG15035 (60404) | 59 | 6 | 0 | 0 | 1.1 | 65 |  |
| Acn + CG17598 (38345) | 24 | 23 | 0 | 24 | 2.3 | 71 | -44.76% |
| CG17598 (38345) | 83 | 5 | 0 | 0 | 1.1 | 88 |  |
| Acn + CG17746 (38347) | 6 | 22 | 0 | 18 | 2.7 | 46 | -14.56% |
| CG17746 (38347) | 74 | 3 | 0 | 0 | 1.03 | 77 |  |
| Acn + CG17746 (105177) | 9 | 16 | 0 | 7 | 2.2 | 32 | -50% |
| CG17746 (105177) | 70 | 3 | 3 | 0 | 1.2 | 73 |  |
| Acn + Fig (65056) | 6 | 8 | 3 | 12 | 2.7 | 39 | -7.55% |
| Fig (65056) | 59 | 4 | 0 | 0 | 1.1 | 63 |  |
| Acn + Mppe (57773) | 6 | 5 | 11 | 25 | 3.2 | 47 | 34.68% |
| Mppe (57773) | 44 | 1 | 0 | 0 | 1.0 | 45 |  |
| Acn + Mppe (5773) | 6 | 5 | 11 | 25 | 3.2 | 47 | 36.36% |
| Mppe (57773) | 44 | 1 | 0 | 0 | 1.1 | 45 |  |
| Acn + Phlpp (57349) | 14 | 16 | 4 | 39 | 3.0 | 73 | 16.51% |
| Phlpp (57349) | 58 | 6 | 0 | 0 | 1.1 | 64 |  |
| Acn + Phlpp (57399) | 14 | 16 | 4 | 39 | 2.9 | 73 | 9.09% |
| Phlpp (57399) | 58 | 6 | 0 | 0 | 1.1 | 64 |  |
| Acn + Pp2C1 (40827) | 10 | 12 | 5 | 38 | 3.1 | 65 | 27.27% |
| Pp2C1 (40827) | 41 | 7 | 0 | 0 | 1.1 | 48 |  |
| Acn + Ppm1 (41987) | 5 | 1 | 2 | 30 | 3.5 | 38 | 33.33% |
| Ppm1 (41987) | 17 | 18 | 20 | 0 | 2.1 | 55 |  |
| Acn + Pgam5 (33346) | 6 | 12 | 5 | 53 | 3.4 | 76 | 30.53% |
| Pgam5 (33346) | 22 | 17 | 21 | 0 | 1.9 | 60 |  |
| Acn + Pgam5-2 (57006) | 6 | 4 | 18 | 12 | 2.8 | 40 | 0% |
| Pgam5-2 (57006) | 44 | 15 | 9 | 0 | 1.5 | 68 |  |
| Acn + Ssu72 (57018) | 0 | 16 | 2 | 11 | 2.8 | 29 | 0% |
| Ssu72 (57018) | 25 | 5 | 0 | 0 | 1.2 | 30 |  |
| Acn + CanA1 (25850) | 19 | 6 | 0 | 45 | 3.0 | 70 | 20.39% |
| CanA1 (25850) | 65 | 2 | 0 | 0 | 1.0 | 67 |  |
| Acn + CG11597 (61988) | 13 | 10 | 0 | 8 | 2.5 | 31 | 25% |
| CG11597 (61988) | 49 | 7 | 0 | 0 | 1.1 | 56 |  |
| Acn + CG11597 (57047) | 18 | 3 | 0 | 24 | 2.9 | 45 | 8.18% |
| CG11597 (57047) | 64 | 6 | 0 | 0 | 1.1 | 70 |  |
| Acn + CG11597 (43175) | 18 | 0 | 0 | 41 | 3.1 | 59 | 18.30% |
| CG11597 (43175) | 45 | 35 | 0 | 0 | 1.5 | 75 |  |
| Acn + Flw (38336) | 24 | 12 | 0 | 45 | 2.8 | 81 | 0.91% |
| Flw (38336) | 65 | 6 | 0 | 0 | 1.1 | 71 |  |
| Acn + Flw (57022) | 5 | 1 | 0 | 24 | 3.6 | 34 | 68.91% |
| Flw (57022) | 39 | 5 | 0 | 0 | 1.1 | 44 |  |
| Acn + Mts (27723) | 0 | 0 | 2 | 64 | 4.0 | 66 | 98.33% |
| Mts (27723) | 37 | 7 | 3 | 0 | 1.2 | 50 |  |
| Acn + Mts (60342) | 11 | 13 | 2 | 9 | 2.3 | 35 | -33.42% |
| Mts (60342) | 15 | 25 | 0 | 0 | 1.6 | 40 |  |
| Acn + Mts (57034) | 0 | 0 | 0 | 2 | 4.0 | 2 | 42.86% |
| Mts (57034) | 0 | 0 | 0 | 1 | 4 | 1 |  |
| Acn + Pp1-13C (32465) | 3 | 4 | 0 | 48 | 3.7 | 55 | 66.92% |
| Pp1-13C (32465) | 30 | 15 | 0 | 0 | 1.3 | 45 |  |
| Acn + Pp1-13C (107770) | 6 | 17 | 13 | 18 | 2.8 | 54 | 0% |
| Pp1-13C (107770) | 55 | 20 | 17 | 5 | 1.8 | 92 |  |
| Acn + Pp1-87B (32414) | 67 | 0 | 0 | 0 | 4.0 | 67 | 33.33% |
| Pp1-87B (32414) | 0 | 0 | 3 | 27 | 3.6 | 30 |  |
| Acn + Pp1α-96A (40906) | 29 | 8 | 0 | 14 | 2.1 | 48 | -61.65% |
| Pp1α-96A (40906) | 27 | 6 | 0 | 0 | 1.1 | 33 | |
| Acn + Pp1α-96A (42641) | 14 | 10 | 22 | 0 | 2.4 | 46 | -22.22% |
| Pp1α-96A (42641) | 10 | 3 | 0 | 6 | 1.8 | 19 | |
| Acn + Pp1-Y1 (58098) | 13 | 13 | 9 | 6 | 2.2 | 41 | -54.95% |
| Pp1-Y1 (58098) | 64 | 8 | 0 | 0 | 1.1 | 72 |  |
| Acn + Pp1-Y1 (65924) | 8 | 2 | 0 | 35 | 3.4 | 45 | 33.33% |
| Pp1-Y1 (65924) | 15 | 18 | 4 | 0 | 1.8 | 37 |  |
| Acn + Pp1-Y2 (57236) | 18 | 10 | 0 | 23 | 2.6 | 51 | -19.69% |
| Pp1-Y2 (57236) | 47 | 18 | 0 | 0 | 1.3 | 65 |  |
| Acn + Pp2B-14D (25929) | 31 | 23 | 6 | 13 | 2.0 | 73 | -72.48% |
| Pp2B-14D (25929) | 48 | 5 | 0 | 0 | 1.1 | 53 |  |
| Acn + Pp4-19C (27726) | 18 | 17 | 3 | 19 | 2.4 | 57 | -36.76% |
| Pp4-19C (27726) | 62 | 6 | 0 | 0 | 1.1 | 68 |  |
| Acn + Pp4-19C (57823) | 0 | 3 | 19 | 54 | 3.3 | 76 | 32.86% |
| Pp4-19C (57823) | 15 | 10 | 27 | 0 | 2.1 | 52 |  |
| Acn + PpD3 (42794) | 8 | 8 | 7 | 17 | 2.8 | 40 | 2.29% |
| PpD3 (42794) | 38 | 4 | 0 | 0 | 1.1 | 42 |  |
| Acn + PpD6 (62849) | 3 | 3 | 1 | 26 | 3.5 | 33 | 48.27% |
| PpD6 (62849) | 28 | 25 | 0 | 0 | 1.5 | 53 |  |
| Acn + PpN58A (57402) | 16 | 20 | 2 | 36 | 2.8 | 74 | -1.28% |
| PpN58A (57402) | 40 | 53 | 0 | 0 | 1.6 | 93 |  |
| Acn + PpV (36064) | 3 | 3 | 2 | 25 | 3.5 | 33 | 48.90% |
| PpV (36064) | 32 | 18 | 2 | 0 | 1.4 | 52 |  |
| Acn + PpV (57765) | 23 | 26 | 6 | 12 | 2.1 | 67 | -56.00% |
| PpV (57765) | 57 | 19 | 0 | 0 | 1.3 | 76 |  |
| Acn + PpY-55A (57519) | 33 | 19 | 0 | 0 | 1.4 | 52 | -113.89% |
| PpY-55A (57519) | 37 | 13 | 0 | 0 | 1.3 | 50 |  |
| Acn + RdgC (60076) | 23 | 28 | 1 | 29 | 2.5 | 71 | -17.44% |
| RdgC (60076) | 14 | 36 | 0 | 0 | 1.7 | 50 |  |

All flies were raised at 28°C.
Scores to calculate Average Roughness: normal = 1; mild = 2; rough = 3; strongly rough = 4.
(1) Positive or negative numbers indicate suppression and enhancement, respectively.
(2) Green or red colors highlight UAS-transgenes with more than 45% suppression or enhancement.
(3) Numbers in parenthesis indicated stock numbers of the Bloomington Drosophila stock center

**Supplemental Table 2.** Effect of loss and gain of phosphatase activity on eye pigmentation in a Drosophila Huntington’s model.

| Genotype | Scores for Depigmented Fly Eye |  |  |  |
| --- | --- | --- | --- | --- |
|  | 1 | 2 | 3 | 4 |
| GMR>Q93 | 0 | 8 | 56 | 39 |
| GMR>Q93+Nil RNAi | 5 | 72 | 17 | 0 |
| GMR>Q93+hPPM1B | 0 | 17 | 45 | 31 |
| GMR/+ | 93 | 0 | 0 | 0 |
| GMR>Nil RNAi | 97 | 0 | 0 | 0 |
| GMR>hPPM1B | 72 | 0 | 0 | 0 |

All flies were raised at 28°C.
Scores to calculate Depigmentation and Roughness
Score 1: No depigmentation and no roughness
Score 2: Mild depigmentation and no roughness
Score 3: Moderate depigmentation and roughness
Score 4: Extreme depigmentation and roughness

**Supplemental Table 3.**
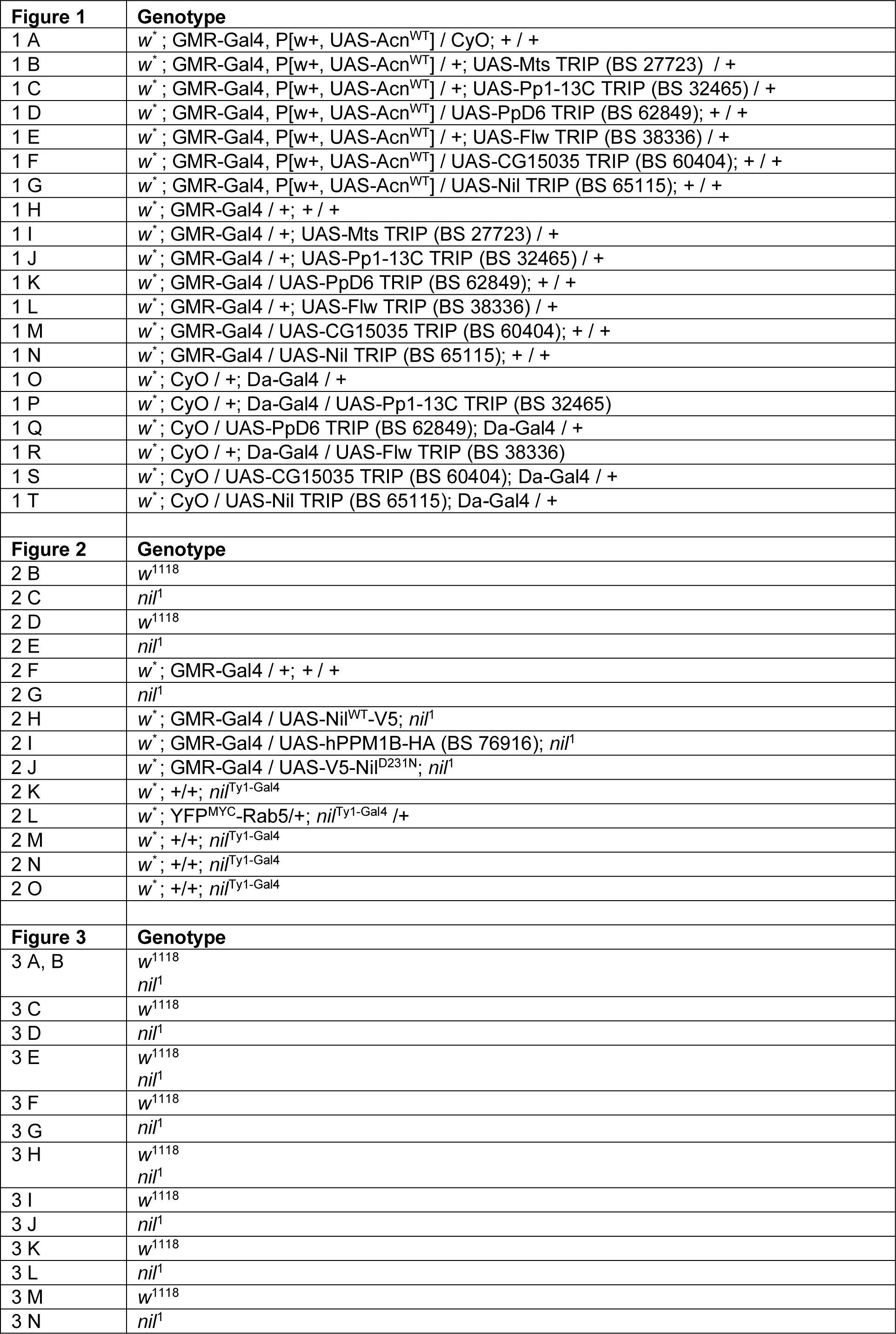
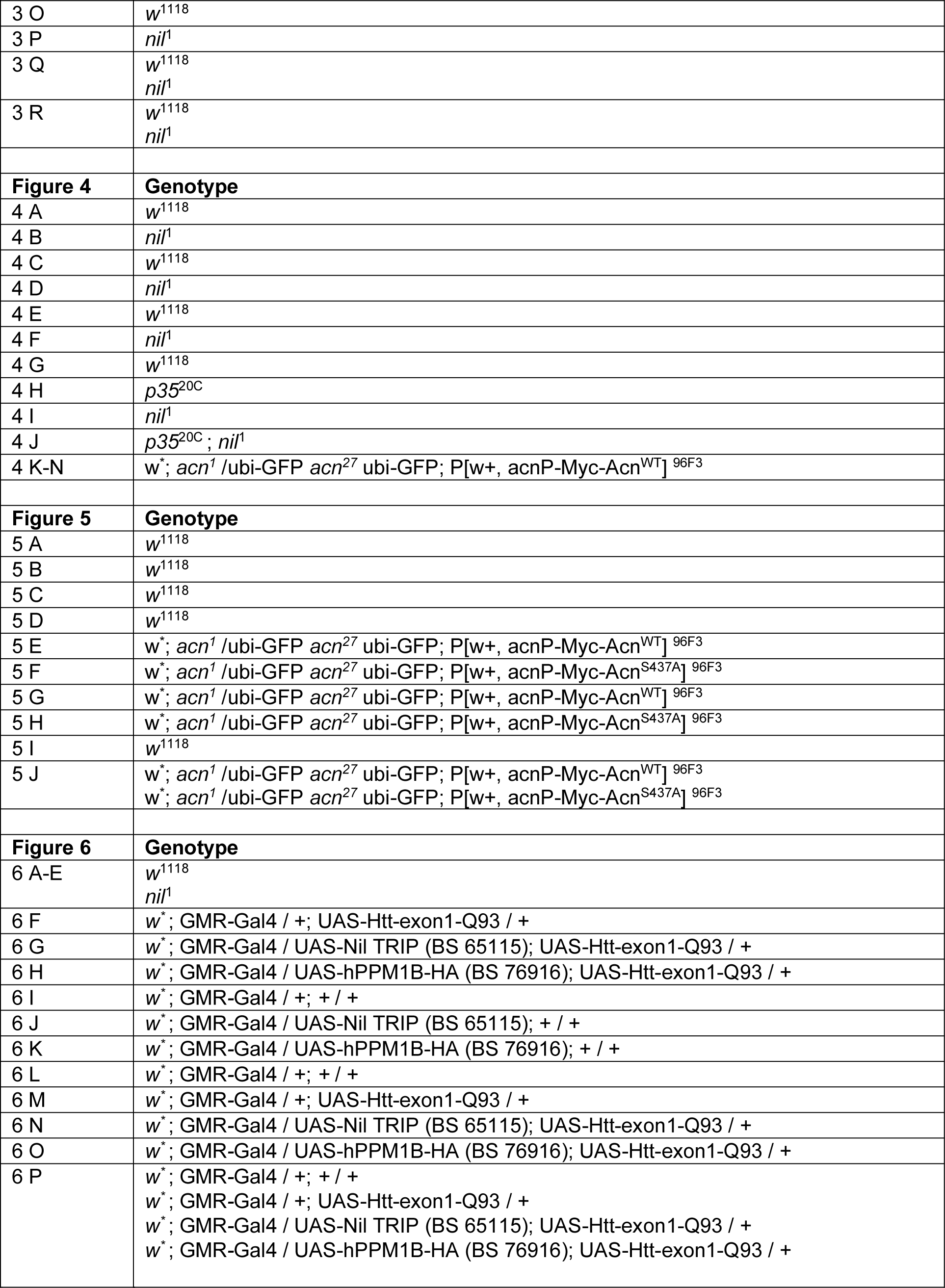
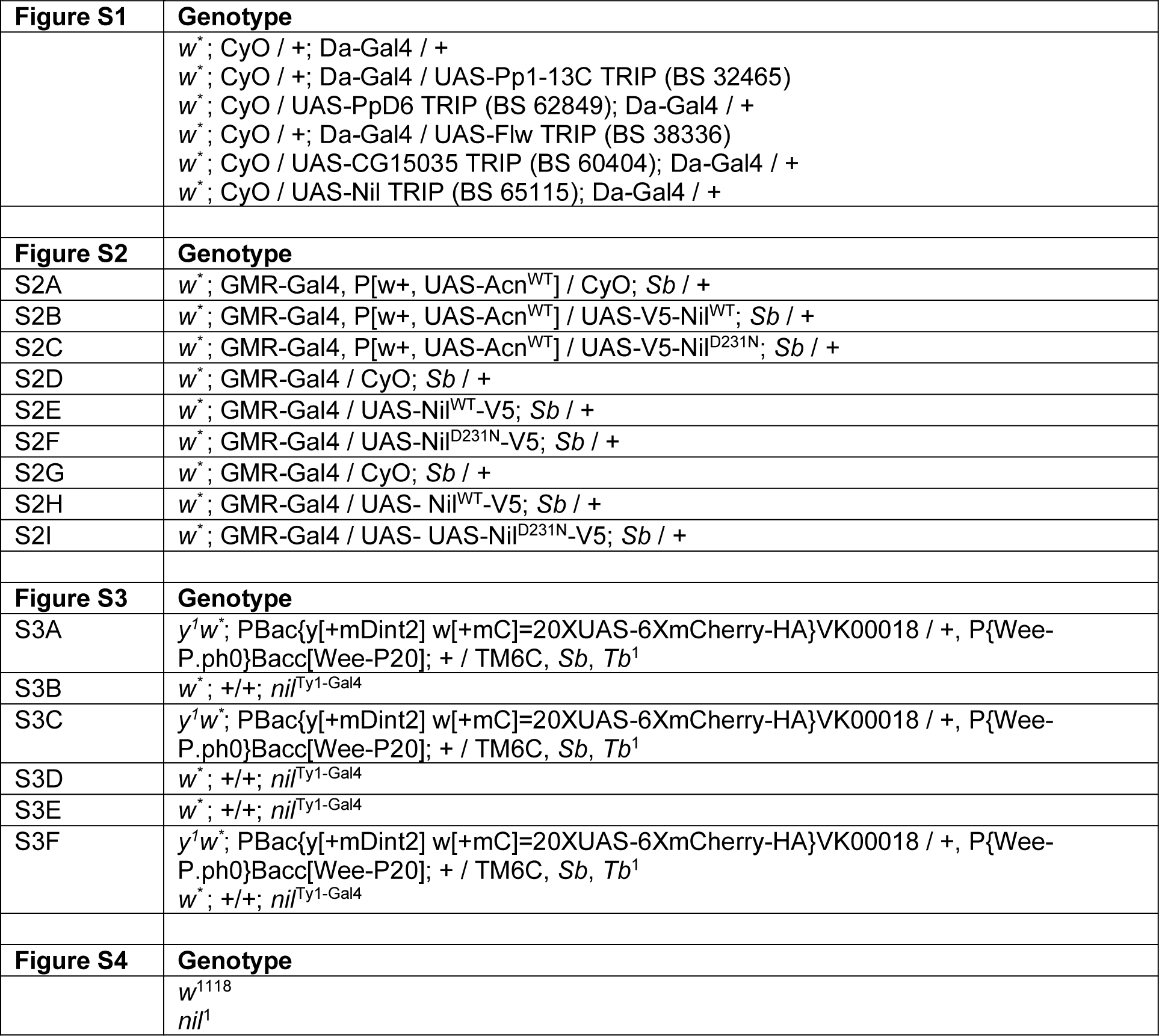
Genotypes of Flies Used for Each Figure

**Supplemental Table 4.**
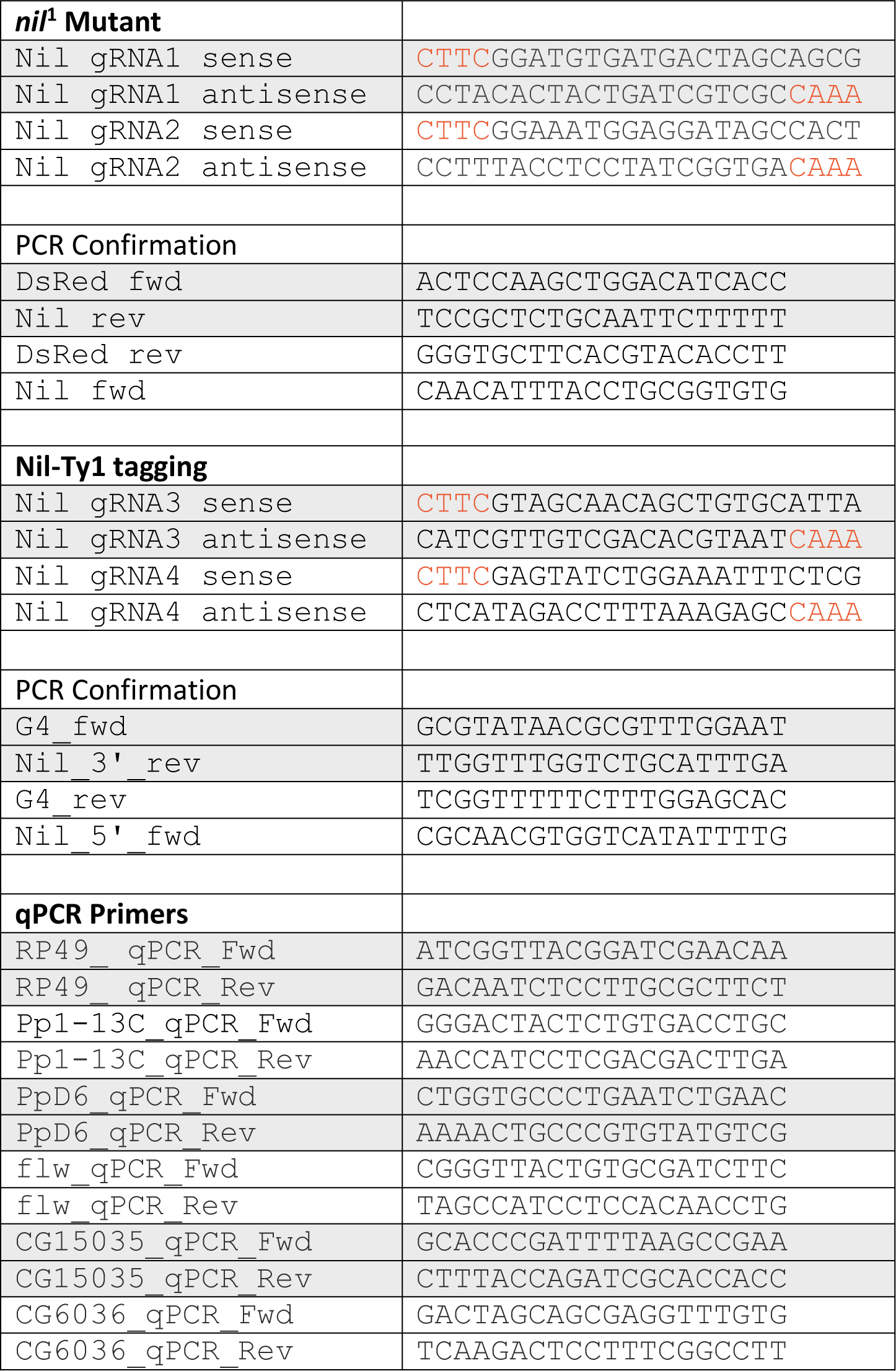
DNA oligonucleotides used

Figure 2- figure supplement 2- source data 1

Raw western blot data with molecular weight markers for Figure 2-figure supplement 2F from lysates of adult male flies of *nil*^Ty1-G4^ and appropriate control probed for Ty1 and Hook.

Western blot analysis using anti-Ty1 and anti-Hook antibodies in lysates from adult males of *nil*^Ty1G4^ and appropriate control. The parts of the raw image used in Figure 2-figure supplement 2F were marked with box.

Figure 3—source data 1

Raw western blot data with molecular weight markers for Figure 3A from lysates of adult heads of *w*^1118^ and *nil*^1^ probed for ATG8a and Actin.

Western blot analysis using anti-ATG8a and anti-actin antibodies in lysates from adult heads of *w*^1118^ and *nil*^1^. Boxes mark the parts of the raw image used in Figure 3A.

